# A robust field-based method to screen heat tolerance in wheat

**DOI:** 10.1101/2021.06.09.447803

**Authors:** Najeeb Ullah, Jack Christopher, Troy Frederiks, Shangyu Ma, Daniel KY Tan, Karine Chenu

**Affiliations:** The University of Queensland, Queensland Alliance for Agriculture and Food Innovation (QAAFI), Leslie Research Facility, 13 Holberton street, Toowoomba, QLD 4350, Australia; Department of Agriculture and Fisheries, 203 Tor street, Toowoomba, QLD 4350, Australia; School of Agronomy, Anhui Agricultural University, Hefei, Anhui, China; The University of Sydney, Plant Breeding Institute, Sydney Institute of Agriculture, School of Life and Environmental Sciences, Faculty of Science, Sydney, NSW 2006, Australia

**Keywords:** Phenotyping, heat stress, genotype x environment interaction, crop improvement, breeding, photoperiod extension

## Abstract

Wheat is highly sensitive to elevated temperatures, particularly during pollen meiosis and early-to-mid grain filling. The impact of heat stress greatly depends on the plant developmental stage. Thus, germplasm ranking for heat tolerance in field trials may be confounded by variations in developmental phase between genotypes at the time of heat events. A photoperiod-extension method (PEM) was developed allowing screening of 35 diverse genotypes at matched developmental phase despite phenological variations. Paired trials were conducted to compare the new PEM against conventional field screening in plots. In the PEM, plants were sown in single rows or small plots. Artificial lighting was installed at one end of each row or plot to extend day length, inducing a gradient of flowering times with distance from the lights. Individual stems or plot quadrats of each genotype were tagged at flowering. Late-sown plants received more heat shocks during early to mid grain filling than earlier sowings, suffering reductions in both individual grain weight (IGW) and yield. IGW was reduced by 1.5 mg for each additional post-flowering day with temperature > 30°C. Significant genotypic differences in heat tolerance ranking were observed between PEM versus conventional plot screening. Strong correlations between trials experiencing similar degree of heat were found both for IGW and for total grain weight with the PEM either with individual-stem tagging (e.g. average *r* of 0.59 and 0.54, respectively for environments with moderate postflowering heat) or quadrat tagging (*r* of 0.53 and 0.47). However, correlations for IGW and yield in these environments were either poor or negative for conventional trials (e.g. average *r* of 0.11 and 0.12, respectively for environments with moderate postflowering heat). Accordingly, a PCA grouped genotypes consistently for heir performance across environments with similar heat stress in PEM trials but not in conventional trials. In this study, most consistent genotype ranking for heat tolerance was achieved with the PEM with tagging and harvesting individual spikes at matched developmental phase. The PEM with quadrat sampling provided slightly less consistent rankings but appears overall more suitable for high-throughput phenotyping. The method promises to improve the efficiency of heat tolerance field screening, particularly when comparing genotypes of different maturity types.

## Introduction

Climate variability is among the major determinants of global crop yields, including wheat (Najeeb et al., 2019; Ray et al., 2015) and strongly impedes plant breeding (Chapman et al., 2012). A significant increase in the frequency of heat stress events, particularly during grain filling has been recorded in the major wheat production regions such as Australia during the past 30 years (Ababaei and Chenu, 2020). These heat events have substantially affected the growth, development and ultimately yield of wheat crops (e.g. Ababaei and Chenu, 2020; Zheng et al., 2016), and are impacting optimal management practices (e.g. Collins and Chenu, 2021; Lobell et al., 2015; Zheng et al., 2012). With the recent rate of climate change, a further increase in the frequency of these heat events is projected in the near future both in Australia (Collins and Chenu, 2021) and globally (Field et al., 2012). Thus, developing wheat genotypes with superior heat tolerance during grain filling is critical for sustaining wheat grain yields and maintaining food security in future hot climates (Ullah et al., 2020).

Wheat is highly sensitive to elevated temperatures and the impact of heat stress is strongly dependent on the crop developmental stage (e.g. Chenu and Oudin, 2019; Farooq et al., 2011; Prasad and Djanaguiraman, 2014). Reproductive and grain filling phases of wheat are extremely sensitive to heat and even a mild increase in the atmospheric temperature during these stages can significantly reduce grain yield. For example, a single hot day (>30°C) during early reproductive stage or onset of meiosis in pollen or micro / megaspore development, can completely sterilise the developing wheat pollen (Saini and Aspinall, 1982). The effect of high temperature on pollen typically translates into a poor grain set and grain yield loss (Guo et al., 2016). Each degree increase (from 15–22°C) in mean temperature during pollen developmental can reduce grain number per unit area by 4%, while a degree increase in maximum temperature at mid anthesis can also result in a 4% reduction in grain number in wheat (Wheeler et al., 1996).

Post-anthesis heat reduces grain yield primarily by limiting assimilate synthesis, translocation, and starch deposition to developing grains (Sofield et al., 1977). Grain weight is most sensitive to heat during early grain filling and becomes progressively less sensitive as grain filling proceeds (Stone and Nicolas, 1998). A single hot day (maximum temperature of 40°C) occurring 10–13 days after anthesis can reduce individual grain weight (IGW) by 14% (Stone and Nicolas, 1998). In addition, IGW may be reduced by 0.5% for each day delay in exposure to heat stress between 15 and 35 days after anthesis (Stone and Nicolas, 1998). Reduction in IGW of heat stressed plants is strongly linked with a shortened grain filling duration (Girousse et al., 2021; Stone and Nicolas, 1998). For each °C rise in mean daily temperature above optimum (15–20°C), a two to eight day reduction in grain filling duration has been reported in wheat crop (reviewed by Streck, 2005). In the Australian wheatbelt, a steady increase in the frequency of hot days (T_max_ > 26°C) during the grain filling period of wheat crops has been recorded over the past 30 years (Ababaei and Chenu, 2020). These authors also pointed out that heat-induced yield losses due to reduced grain weight (18.1%) were greater than those due to grain number (3.6%). This highlights the importance of developing wheat germplasm more tolerant to heat post-flowering.

Conventionally, wheat genotypes are screened for heat tolerance by serial sowings, using heat chambers in the field, or in controlled environments (e.g. Telfer et al., 2021, 2018; Thistlethwaite et al., 2020). Ranking for heat tolerance is typically based on physiological or morphological traits associated with plant function and performance (Bennett et al., 2012). However, changes in these traits are strongly influenced by the environment and the methodology used (Limpens et al., 2012; Poorter et al., 2016). Field-based screening methods are generally considered more representative of the plant response to natural environments (Passioura, 2006). However, screening of wheat genotypes with varying maturity types may be complicated by the unpredictability of heat events under field conditions. The impact of heat events is highly dependent on the developmental phase (e.g. Chenu and Oudin, 2019; Djanaguiraman et al., 2014; Tashiro and Wardlaw, 1990), which is specific to each particular wheat line being tested. Thus, ranking of wheat genotypes for heat tolerance may be confounded by variation in developmental stage during a natural heat event. An improved technique to screen for high temperature stress at matched developmental stages of wheat genotypes in the field could accelerate the selection of heat tolerant genotypes.

Here, we developed and tested a photoperiod-extension method (PEM) that allows comparison of the performance of wheat genotypes with varying maturity types at common developmental stages during natural heat events. The method was tested across three different locations over three consecutive years, harvesting either specific spikes that flowered at the same time or small areas of plants (i.e. quadrats) flowering the same day (i.e. 50% of the spikes at anthesis). Rankings of wheat genotypes with varying maturity type and heat tolerance were compared using PEM and conventional plots.

## Materials and Methods

### Growth conditions and experimental design

Field trials were conducted over three consecutive years (2018 to 2020) at three locations across southern Queensland, Australia. Conventional plots were estabilished adjacent to trials using the newly developed photoperiod-extension method (PEM). Randomised block design trials with two times of sowing blocks and four replicates were established each year. To maximise the likelihood of heat during grain filling, trials were planted typically later in the cropping season than industry practice (Table 1). At The University of Queensland Research Farm, Gatton (27°34′50″S, 152°19′28″E), the first sowings (s1) were established between late May or early July and the second sowings (s2) between late August or early September. At the Hermitage Research Station, Warwick (28°12’40’’S, 152°06’06’’E), trials were sown early June or mid-July (s1) or mid-August and mid-September (s2). At the Tosari Crop Research Centre, Tummaville (27°49’09.1”S 151°26’14.9”E), plants were sown mid-July (s1) and early September (s2). All trials were fully irrigated (except at Tosari) and cultivated under non-limiting fertiliser conditions (Table 1). A boom irrigator (centre pivot sprinkler) was used for irrigating plots at Gatton (2018-2020) as well as both plot and PEM trials at Tosari. PEM trials at Gatton (2018-2020) and Warwick (2018) were irrigated with wobbler sprinklers. In 2020, both PEM and conventional plots trials at Warwick were irrigated using a drip irrigation system. At Tosari, trials were irrigated at sowing and pre-flowering, but no post-flowering irrigation was applied, as water was then unavailable. For all trials, standard crop management practices including weed, disease and pest control were adopted during the season.

**Table 1.** Trial characteristics, including the trial identifier (Trial), site, irrigation treatment, sowing date and the types of trial conducted (i.e. PEM with tagging and harvesting of either single spikes or quadrates; and conventional plots). Also presented are mean and max daily temperature, day-time vapour pressure deficit (VPD) and radiation from sowing to maturity, as well as the mean duration of the pre- and post-flowering periods, and the heat environment type (HET) for the first tagging of trials with the photoperiod-extension method (PEM). Days to flowering and post-flowering duration were calculated from sowing to flowering and flowering to maturity, respectively. TOS19 trial only had supplementary pre-flowering irrigation and experienced a mild post-flowering water stress.

| Trial | Site | Sowing | Trial type | Sowing date | Irrigation | Mean temp. (°C) | Mean daily max temp. (°C) | Mean VPD (kPa) | Mean daily radiation (MJ m<sup>-2</sup>) | Days to flowering (days) | Post-flowering duration (days) | HET |
| --- | --- | --- | --- | --- | --- | --- | --- | --- | --- | --- | --- | --- |
| GAT18-s1* | Gatton | s1 | Spike & quadrate PEM | 03/07/2018 | Full | 19.3 | 26.3 | 0.74 | 23.8 | 73 | 42 | HET1 |
| GAT18-s2 | Gatton | s2 | Spike & quadrate PEM & plots | 31/08/2018 | Full | 23.1 | 31.4 | 1.38 | 23.2 | 53 | 39 | HET2 |
| WAR18-s1 | Warwick | s1 | Spike & quadrate PEM | 16/07/2018 | Full | 18.6 | 25.6 | 0.84 | 17.7 | 73 | 39 | HET1 |
| WAR18-s2 | Warwick | s2 | Spike & quadrate PEM | 12/09/2018 | Full | 22.0 | 30.3 | 1.52 | 21.2 | 52 | 37 | HET2 |
| GAT19-s1 | Gatton | s1 | Spike PEM & plots | 09/07/2019 | Full | 19.5 | 28.7 | 1.40 | 16.8 | 71 | 39 | HET2 |
| GAT19-s2 | Gatton | s2 | Spike PEM & plots | 03/09/2019 | Full | 23.7 | 33.8 | 2.23 | 20.8 | 59 | 35 | HET3 |
| TOS19-s1 | Tosari | s1 | Spike & quadrate PEM & plots | 16/07/2019 | Supplementary | 21.1 | 30.2 | 1.26 | 17.7 | 79 | 35 | HET2 |
| TOS19-s2 | Tosari | s2 | Spike PEM & plots | 06/09/2019 | Supplementary | 26.5 | 36.7 | 2.27 | 16.2 | 62 | 35 | HET3 |
| GAT20-s1 | Gatton | s1 | Quadrate PEM & plots | 26/05/2020 | Full | 17.9 | 26.8 | 1.08 | 17.3 | 79 | 49 | HET1 |
| GAT20-s2 | Gatton | s2 | Quadrate PEM & plots | 04/08/ 2020 | Full | 22.2 | 31.2 | 1.35 | 19.7 | 65 | 37 | HET2 |
| WAR20-s1 | Warwick | s1 | Quadrate PEM & plots | 08/06/ 2020 | Full | 19.3 | 26.3 | 0.74 | 13.6 | 73 | 42 | HET1 |
| WAR20-s2 | Warwick | s2 | Quadrate PEM & plots | 12/08/2020 | Full | 23.1 | 31.4 | 1.38 | 11.6 | 53 | 39 | HET1 |

Trial identifiers indicate site (Gatton (GAT), Tosari (TOS) or Warwick (WAR)), year conducted and sowing time (s1 or s2).

With the PEM, each wheat genotypes was either hand sown in a 5 m single row (33 cm row spacing) in 2018 and 2019, or machine planted in a four row plot (1×5 m, 2020 with 25 cm row spacing).

Conventional field plots were planted at the same time and with similar management to the PEM trials. In 2018 and 2020, the conventional field plots (2×6 m) were set up in Gatton and Warwick, while in 2019, genotypes were tested in smaller plots (1×6 m) at Tosari and Gatton. All conventional plots were planted at a 25 cm row spacing with a population density of 130 plants m^−2^ and in four independent replications. Due to unavailability of irrigation water, only sowing 2 (s2) plots were established at Gatton during 2018.

Trials are given identifiers denoting the site (Gatton (GAT), Tosari (TOS) or Warwick (WAR), and the year of the trial. Data may also be identified by time of sowing (s1 or s2) and the tagging event (T1, T2, T3) when applicable.

### Genotypes

Thirty-five wheat (*Triticum aestivum* L.) genotypes with contrasting phenology and adaptation were used in the study (Table S1, Supplementary). These included high-performing spring cultivars Suntop, Spitfire, Gregory, Janz, Hartog, EGA Wylie, Corack, Yitpi, Mace and Scout widely cultivated in major cropping regions of Australia. A set of eight CIMMYT genotypes described as heat tolerant under Australian environments was obtained from the University of Sydney (Thistlethwaite et al., 2020). Other genotypes used in these trials included donors of a multi-reference parent nested association mapping (MR-NAM) population developed for screening for heat and drought tolerance in wheat (Christopher et al., 2015, 2021; Richard, 2017). In total, 35 wheat genotypes were tested in PEM and plot trials, with 32 genotypes at each trial, except for the PEM trial with quadrat harvest in 2020 when only 20 selected genotypes were used (Table S1, Supplementary). In conventional plot trials, 32 genotypes were used in each trial.

### The photoperiod-extension method

The novel photoperiod-extension method (PEM) was based on a method described by Frederiks et al. (2012) originally developed to test for frost damage. At one end of each row or plot, LED lamps (CLA LT401, 9W T40 LED LAMP, 3000K 760LM) with a lumen efficiency ≥ 80, were set up approximately 1 m above the ground level and at spacing of 0.8 m. These lamps supplement light by extending the day length to 20 h (Fig. 1). The intensity of light diminishes with the square of the distance from the lights along the test row, with maximum effect closest to the lights and minimum or no impact at the other end of the row. This variation in light intensity across the rows induced a gradient of flowering times within each row with the plants closest to the light developing more rapidly.

**Figure 1:**
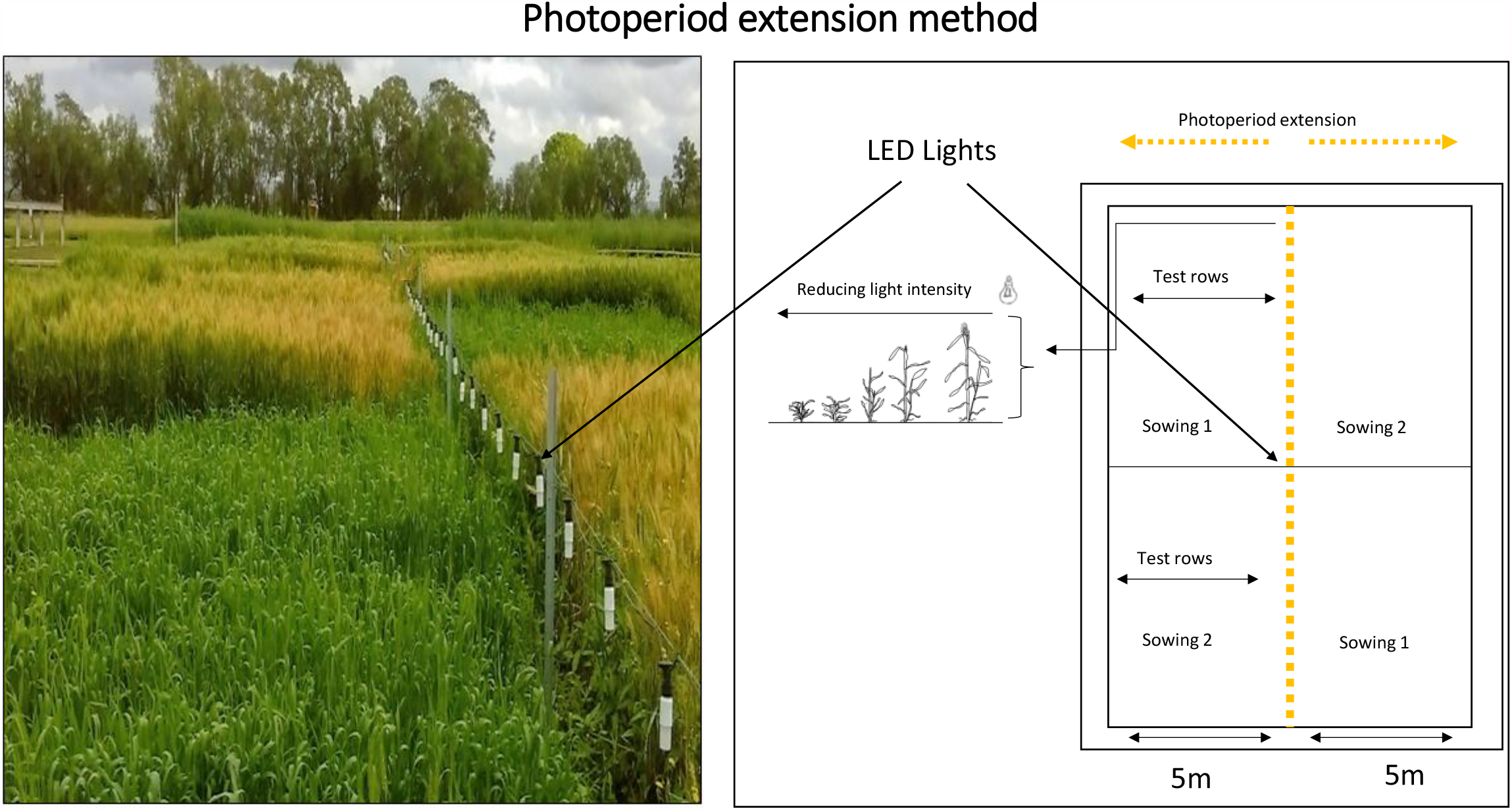
Layout of the photoperiod-extension method (PEM). Wheat genotypes were sown either in a single row (as in this picutre) or in narrow plots of four rows. In the centre of the trial at the end of each test row, at the central axis of the trial, LED lights were setup. These supplemental lights extend the photoperiod to 20 h. The intensity of light diminishes along the row and induces a gradient of flowering times within each test row, with plants closest to the light developing faster.

### Plant measurements

For each PEM trial and sowing date in 2018 and 2019, approximately 20 stems of each genotypes were tagged at flowering (Zadoks decimal growth stage 65; Zadoks et al., 1974). The induced gradient in phenology along the rows allowed tagging of genotypes multiple times for plants in rows or plots from each sowing time. One-to-three cohorts of stems were tagged at precisely matched flowering in rows or plots from each sowing time and trial. These sequentially tagged coherts were termed as ‘tagging 1’ (T1), ‘tagging 2’ (T2) and ‘tagging 3’ (T3). The spikes from tagged stems were manually harvested at maturity and processed for grain yield components.

In addition, in each test row of the all PEM trials in 2018 and s1 trials at Tosari in 2019, a quadrat consisting of a 0.5 m section of row that originally contained heads from tagging 1 was manually harvested to estimate yield and its components. In 2020, ∼ 0.5 m section of each plot (four rows) was tagged at flowering (Zadoks 65), and a quadrat of two central rows (0.5 m each) within the tagged region was manually harvested at crop maturity.

All plots from the conventional method were harvested using a small plot machine harvester at maturity when grain moisture was approximately 11%. Grain samples were maunally counted to calculate individual grain weight (IGW).

### Environment variables

Local weather stations (Campbell Scientific) were set up at each site to record weather data for each 10 min period. Light interception was measured with light sensors (Apogee SP-110 pyranometers, and Apogee SQ-110 for radiation and PAR measurements, respectively) installed at 1.5 m height. HMP60 (Vaisala INTERCAP®) probes were used to measure air temperature (Tair) and relative humidty (RH) at 1.5 m above the ground. Environmental characteristics of each trial are presented in Table 1.

Thermal time was calculated in degree days (°Cd) using the following equation: (Jamieson et al., 1995)

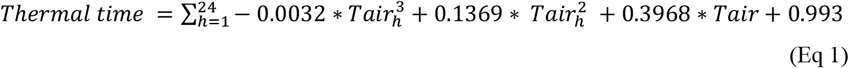

Where Tair_h_ is the hourly air temperature data.

Vapour pressure deficit (VPD) was calculated hourly during the day time as in Alduchov and Eskridge (1996) by the following equation:

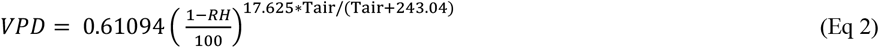

Where Tair and RH were the hourly air temperature and hourly air relative humidity, respectively, during the daytime.

### Statistical analysis

Data were analysed using R (R Core Team, 2018). Individual and interaction effects of genotype and environments (sowing, location and tagging) were determined by analysis of variance (ANOVA). Statistical differences were tested with student’s t-tests at a 5% level. Principle component analysis was computed to rank genotypes in different environments.

## Results

### Wheat crops experienced a wide range of heat events across locations and sowing times

Wheat genotypes at each location, season, sowing and tagging experienced varying air temperatures and vapour pressure deficit (VPD) in the pre- and post-flowering periods (Table 1, Fig. 2 & Table S2). In all trials, plants from sowing 2 (‘s2’) experienced significantly higher temperatures and VPD than those from the first sowing (‘s1’). Post-flowering mean air temperature was ∼20% higher in s2 than s1 across all trials (Table 1). Similar differences in post-flowering daily-maximum air temperature were also observed between s1 and s2, with the exception of in the 2020 Warwick trial (WAR20), where temperatures were only 10% higher. The number of hot days (with maximum temperature >30°C) across these trials also varied greatly, with TOS19s2 in 2019 receiving the maximum number (22) of post-flowering hot days. Higher temperature during s2 shortened the duration of both pre- and post-flowering periods, although the reduction in time to flowering was relatively greater than the reduction in post-flowering duration across all the tested locations (Table 1).

**Figure 2:**
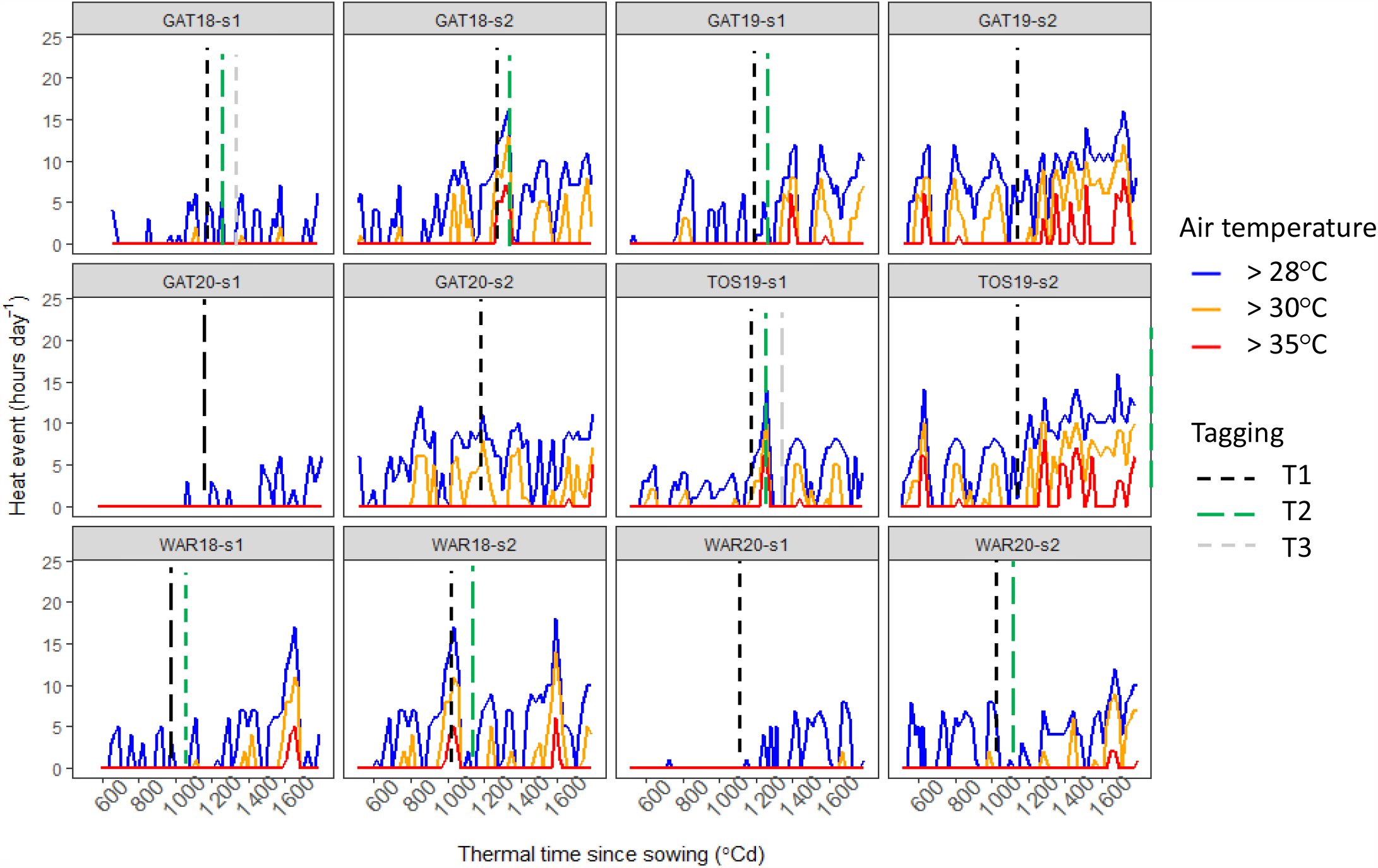
The number of hours each day when air temperatures exceeded 26°C (blue lines), 30°C (orange lines) or 35°C (red lines) is plotted against thermal time (°Cd) from sowing. These are presented for each location, season and sowing time. The vertical lines represent tagging of wheat genotypes at flowering. T1, 1^st^ cohort of stems tagged at flowering, T2: 2^nd^ cohort of stems tagged at flowering, T3: 3^rd^ cohort of stems tagged at flowering. Trial identifiers are as described in Table 1.

Environments with similar post-flowering stress were defined as (i) heat environment type 1 (HET1) that corresponded to environments with no or only late grain-filling heat stress (i.e. less of 4 cumulated hours of temperature >30°C between 0 and 450°Cd after flowering), (ii) HET2 that included all environments with moderate heat stress during grain fill (5 to 15 days with a maximum temperature >30°C in our set of trials), and (iii) HET3 that corresponded to environments severely stressed during grain fill (20 and 22 days with a maximum temperature >30°C in our set of trials). In addition to post-flowering heat, HET3 trials experienced high pre-flowering temperatures (Table S2, Supplementary) including with hot days around stem elongation and meiosis (Fig. 2) that significantly reduced grain set (Table S3 Supplementary).

### Late-sown crops had lower individual grain weight and grain yield

Individual grain weight and grain yield varied significantly across trials, but plants typically produced significantly smaller grains and lower grain yield in s2 compared with s1 (Fig. 3). Trial-mean grain number in s2 was also significantly lower or not significantly different than s1 in almost all trials (Table S3 Supplementary), probably due to shorter pre-flowering periods associated to warmer temperatures (Table S2 Supplementary).

**Figure 3:**
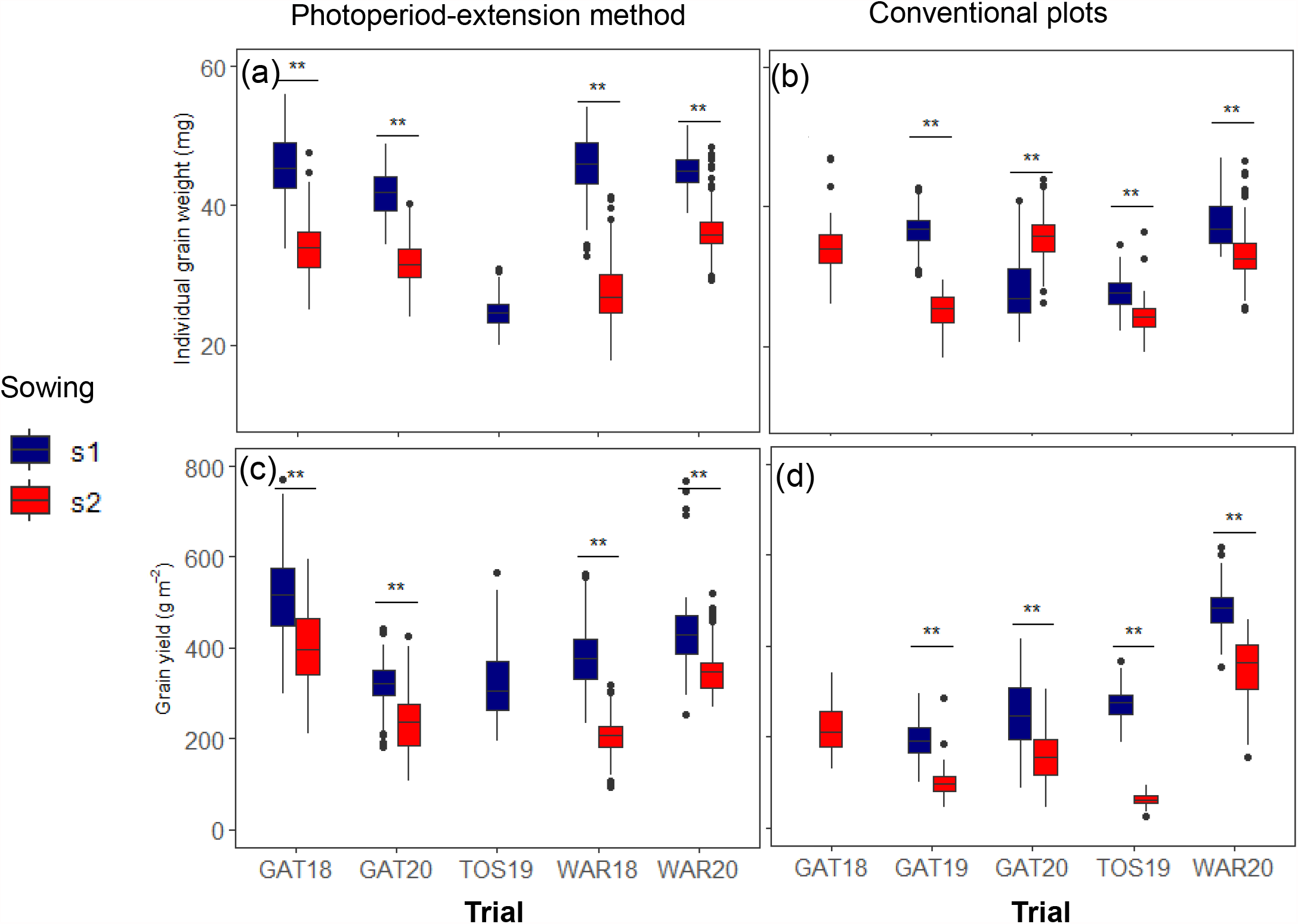
Effect of sowing time on (a, c) individual grain weight and (b, d) grain yield of tested wheat genotypes for trials with (b, d) conventional pots or (a, b) the photoperiod-extension method with quadrat tagging and harvests. Each boxplot displays data from 32 genotypes and four independent replicates. TOS19 crops only had supplementary irrigation and experienced a mild post-flowering water stress. In the boxplot, horizontal black lines insides each box denote median values; boxes extend from the 25^th^ to the 75^th^ percentile of each group’s distribution of values; the whiskers the 10^th^ and 90^th^ percentiles; and the dots outside the whiskers represent individual values outside this range. ** corresponds to significant differences at *P*< 0.001 between the two sowing times within each trial. In 2018, s2 plots were established only at Gatton (GAT18) and in 2019, quadrat harvests were taken from the s1 PEM trials at Tosari (TOS19) only. Trial identifiers are as described in Table 1.

The maximum trial-means for IGW and grain yield were measured in HET1 environments. In the PEM with quadrat tagging and harvests, a maximum of 45.5 mg for IGW and 571 g m^-2^ for grain yield was recorded in GAT18s1 (Fig. 3). With the conventional plot method, genotypes produced a maximum trial-mean IGW of 37.7 mg and a maximum grain yield of 484 g m^-2^ in WAR20s1. Trial-mean reductions in IGW between s2 and s1 for PEM and conventional plot trials were maximum at WAR18 (64%) and GAT19 (47%), respectively (Fig 3a & b). Grain yield reduction between s2 and s1 was maximum in GAT18 for the PEM and in TOS19 for conventional plots (Fig 3c & d).

All trials were fully irrigated, except TOS19, which only received pre-flowering supplementary irrigation and was subjected to a post-flowering drought. Given the pre-flowering conditions, s1 plants at TOS19 produced a high number of grains but significant reduction in grain size was confounded by post-flowering heat and drought.

### Extending the photoperiod increased allowed tagging of plants at a matched development stage from genotypes contrasting maturity

Phenology data were collected from plants 0.5 m from each end of experimental rows in the PEM, from next to the supplemented light and at the far end away from the supplemental lights (Fig. 4-5). The phenology of the different genotypes significantly varied both under natural and supplemented light. Across all PEM trials, the earliest maturing genotypes flowered approximately 10.1 days (172°Cd) and 7.8 days (145°Cd) earlier than the latest maturing genotypes under natural and supplemented lights (Fig. 4). The supplemented light accelerated flowering by 7.9 days (152°Cd) and 5.4 days (114°Cd) on average in s1 and s2, respectively (averaged across genotypes and trials). This gap between flowering times of plants under natural and supplemented light allowed multiple tagging of the genotypes at a matched developmental phase (Zadok 65) from a single time of sowing.

**Figure 4:**
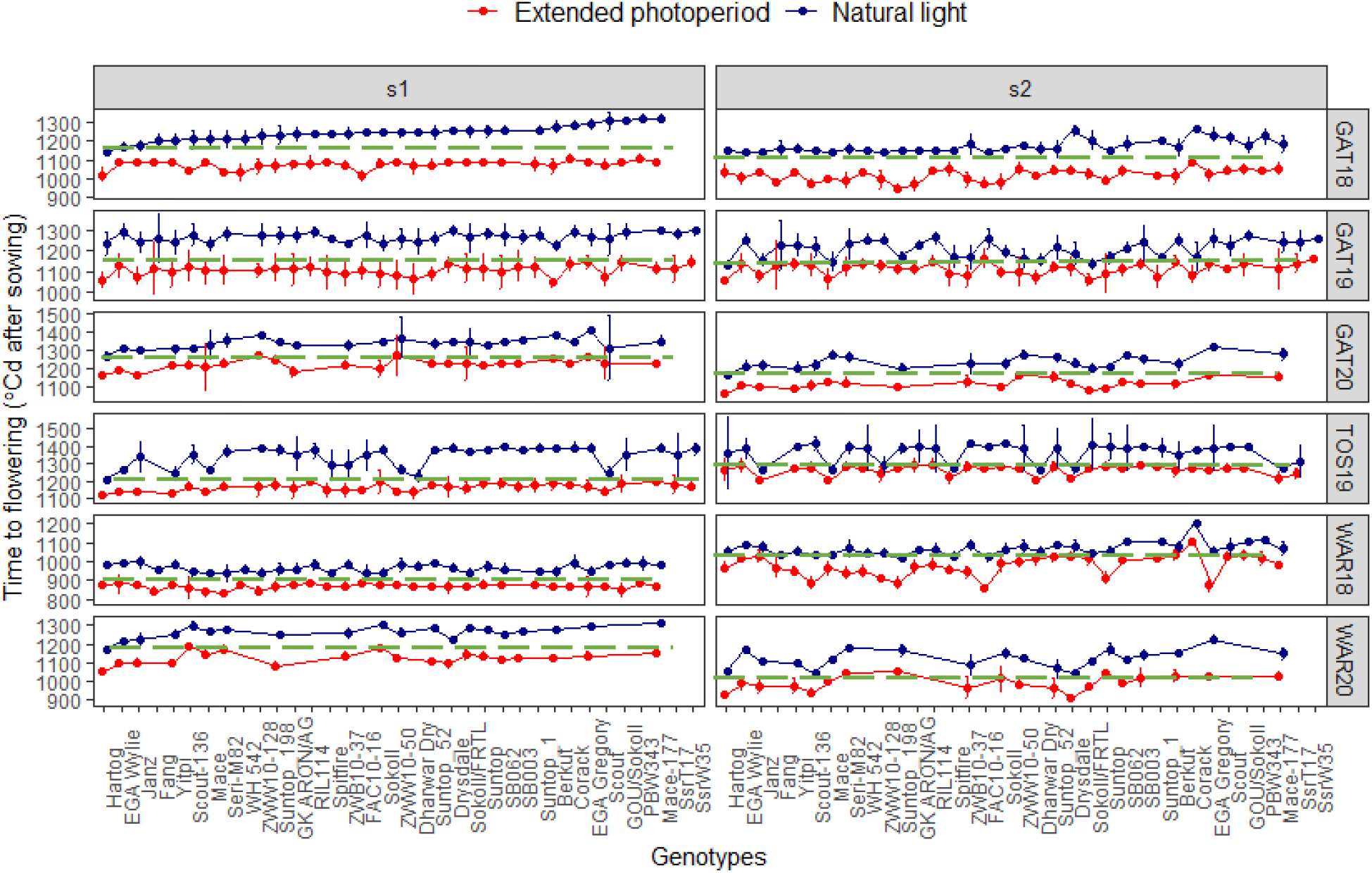
Flowering time of wheat genotypes with and without supplemented light. Horizontal dashed lines correspond to the day of tagging of all wheat genotypes at matched flowering. Data on flowering were collected for plants 0.5 m away from the light source (extended photoperiod, red) and plants 0.5 m from the end of the row (natural light, blue). Values correspond to the mean of four independent replicates ± confidence interval (95%). s1 and s2 represent sowing 1 and sowing 2, respectively.

**Figure 5:**
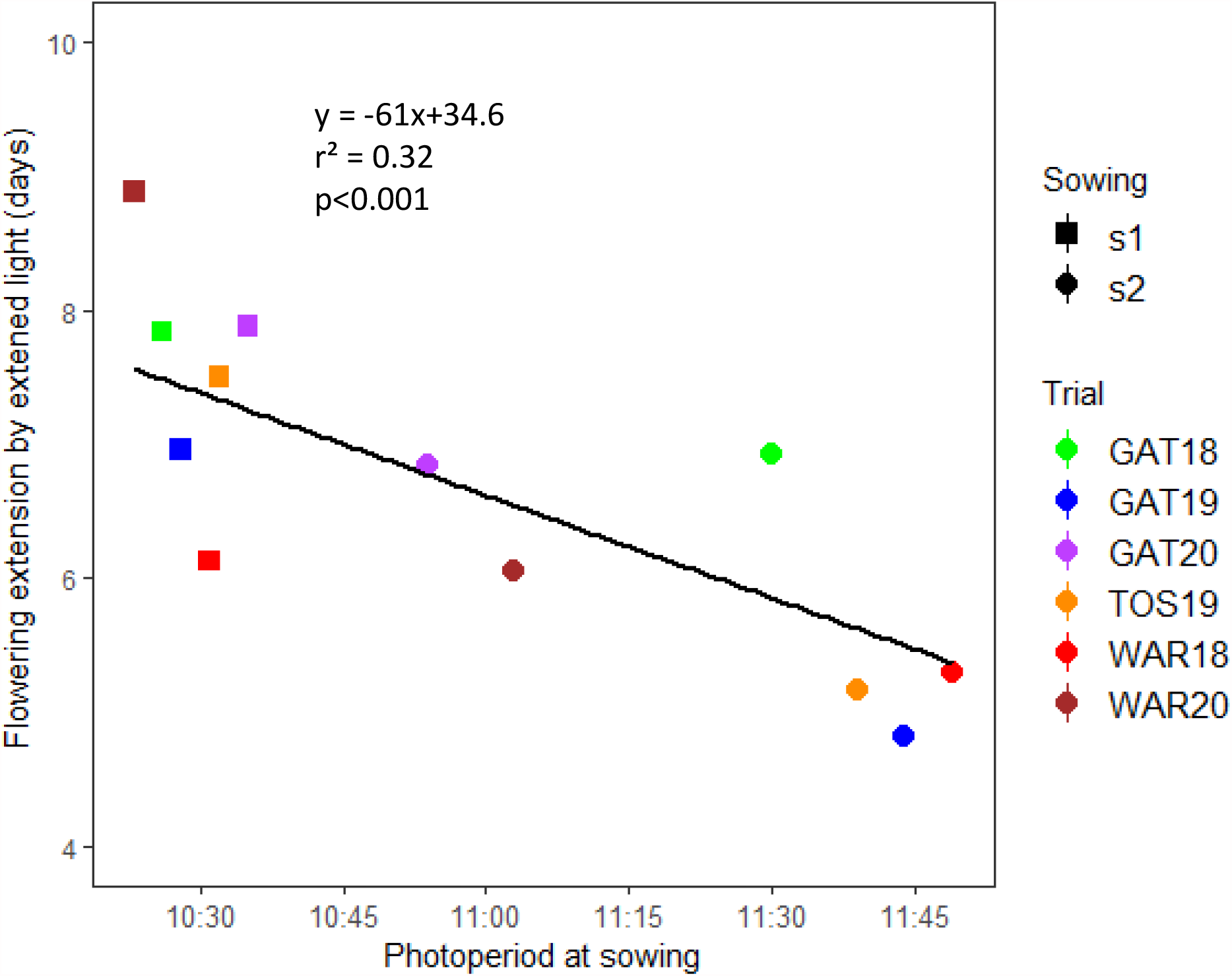
Delay in flowering time due to supplemented light in response to the photoperiod measured at sowing in all trials and sowings. The delay in flowering was calculated as the difference in flowering dates for plants located 0.5 m (i.e. extended light, 20 h) and 4.5 m (i.e. natural light) away from the lights. Data correspond to the mean of 32 (2018 and 2019) or 20 (2020) genotypes and four independent replicates for all PEM plots. TOS19 crops only had supplementary irrigation and experienced a mild post-flowering water stress.

Supplemented light had a weaker effect in late sowings when the natural photoperiod was already longer (Fig. 5). Under the shortest studied photoperiods (<10.5 h at sowing), plants closer to the lights flowered approximately 8.8 days earlier than plants away from the lights (averaged across genotypes and trials). The phenology effect of supplemented light was reduced by approximately 49% when plants were planted under longer photoperiod (11.5 h or more) at sowing.

### A reduction in individual grain weight by 1.5 mg for every post-flowering heat day

A strong correlation (*r*^2^ = 0.90) between the number of post-flowering days with maximum temperature >30°C and individual grain weight was observed across PEM trials with individual-spike harvests (Fig. 6a). Across all trials and tagging events, IGW was reduced by 1.5 mg for every additional post-flowering hot day (with maximum temperature >30°C). Stems receiving 0-4 post-flowering hot days (HET1), produced the largest grains (40-45 mg; Fig. 6a). Given that any heat event affecting these plants were very brief (Fig. 2), IGW produced during this period is likely to be close to the potential grain weight. Stems with a moderate numbers of heat events between 5 and 15 post-flowering hot days (HET2) had significantly lower mean IGW and total grain weight per spike than those in HET1 (Fig. 6). The most stressed stems experienced greater than 15 hot days (HET3, Fig. 3). In GAT19s2 and TOS19s2, most plants experienced 20 and 22 post-flowering hot days, respectively and suffered maximum reduction in IGW. On average, stems under these extremely hot conditions produced 2.7 times smaller grains than the potential IGW (Fig. 6a).

**Figure 6:**
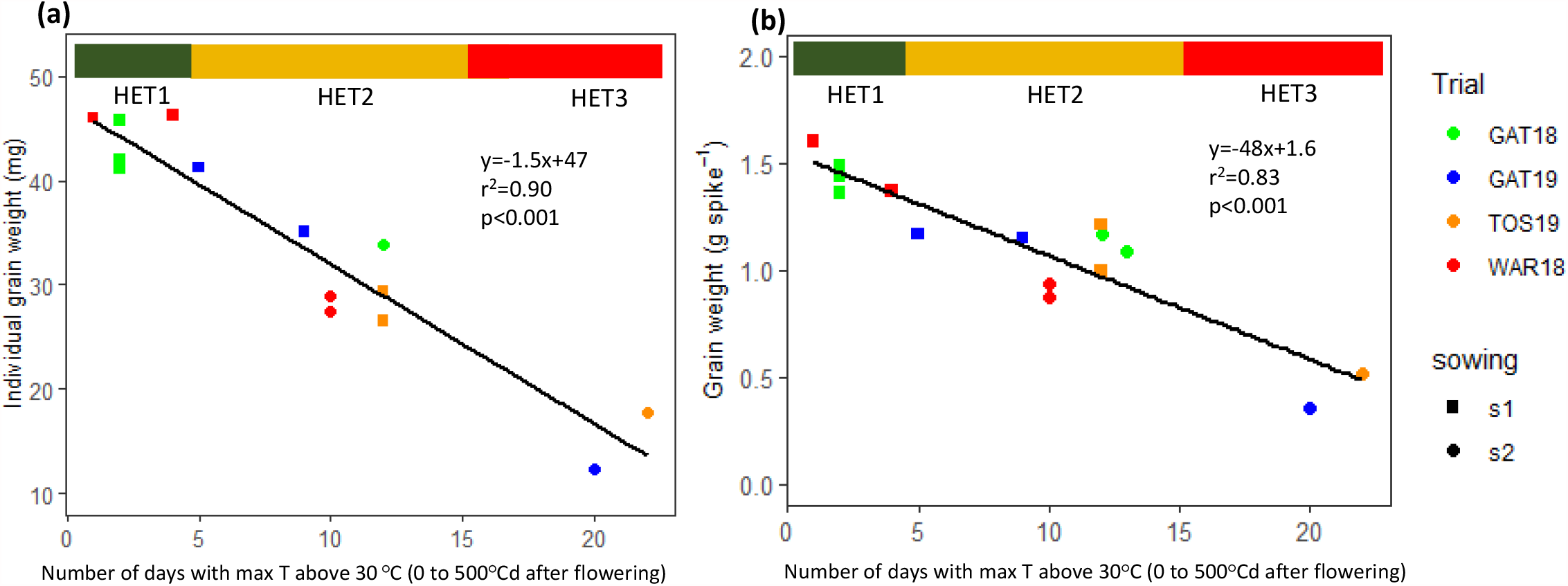
Changes in (a) individual grain weight and (b) spike grain weight, in response to the number of post-flowering hot days (0-500°Cd after flowering) across locations and sowing times. Each point represents the mean of all genotypes for individual grain weight of spikes that flowered the same day (four independent replicates of 20 stems each). Heat environment types are indicated by the horizontal bar at the top of each panel (HET1, green; HET2, orange; HET3, red). TOS19 only had supplementary irrigation and experienced a mild post-flowering water stress.

While greatest environmental variations occurred between sites and sowings, variations in the number of hot days and mean IGW were also observed among stems tagged at different times (i.e. group of stems flowering at different dates) within a single sowing time. For instance, T2 stems of GAT19s1 produced 14% smaller grains compared with T1 stems (Fig. 6a, Table S3).

### The photoperiod-extension method provided a stable genotype ranking within heat environment types

Genotypic rankings for IGW assessed using the PEM were relatively consistent between environments within heat environment types, i.e. environments experiencing similar heat stress (Fig. 7). For single spike harvests with the PEM, IGW correlations for stems experiencing mild or no heat stress during grain-filling (HET1), ranged from *r* of 0.64 to 0.89 (Fig. 7a). In HET2, *r* correlations ranged from 0.2 to 0.90 for fully-irrigated trials (i.e. all trials except TOS19 which experienced a post-flowering drought). These correlations were stronger (0.57-0.9) among the irrigated trials with a similar number of hot days (i.e. 9-13) and became more variable with the environment (GAT19s1T1) with five hot days only (*r* of 0.2-0.57). IGW of the two severely-stressed environments (HET3) were strongly correlated (*r* = 0.46, Fig 7a). Correlations between HET1, HET2 and HET3 varied, with moderate to high correlations between HET1 and HET2 environments and weaker correlations with HET3 environments. IGW from PEM quadrat harvests were also positively correlated between the different tested environments, and these correlations were particularly strong among environments experiencing a similar number of hot days (Fig. 7b). For example, in HET1, these correlations were moderate to strong (*r* of 0.24-0.88) mainly because of a weaker correlation between GAT20s1 and WAR20s1 (*r* = 0.24). Similarly, all fully-irrigated HET2 environments were positively correlated with correlations ranging from 0.19 to 0.68.

**Figure 7:**
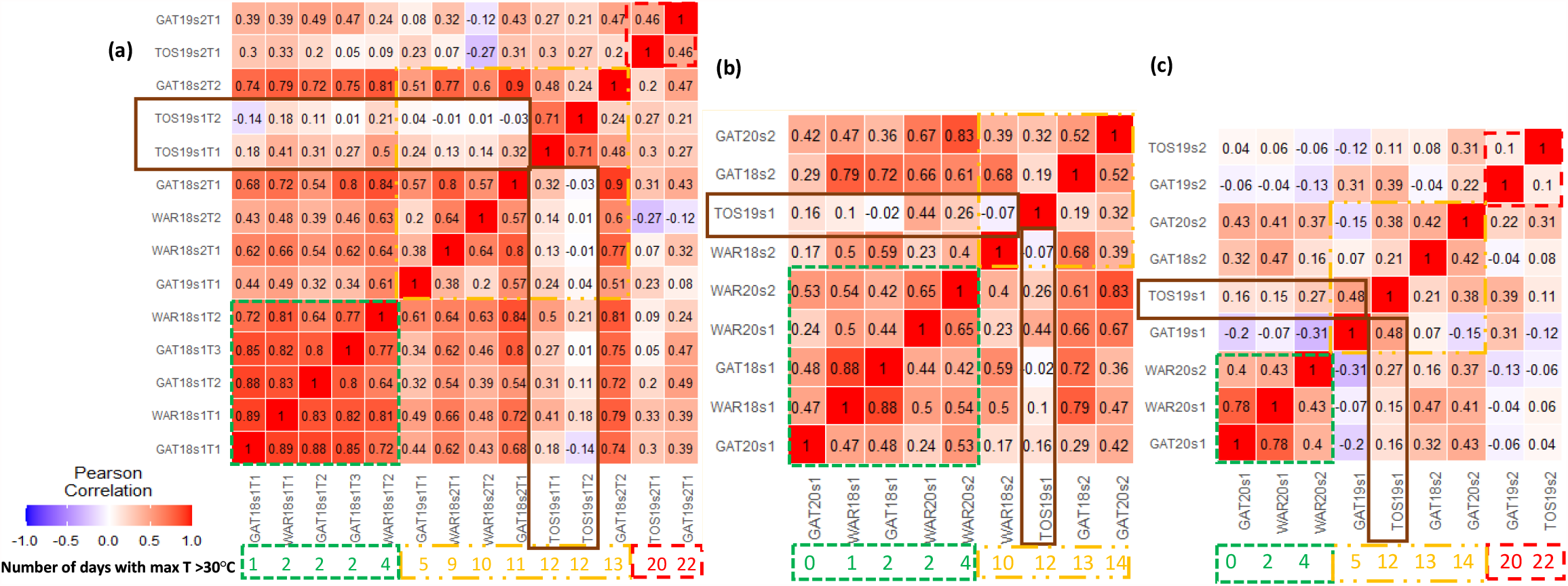
Genetic Pearson’s correlation coefficients (*r*) for individual grain weight between environments. Correlations for mean individual grain weight of genotypes between each pair of tested environments (i.e. site x year x sowing x tagging-event combinations). Individual grain weight was estimated from measurements at (a) spike level, and at (b) crop level with quadrat harvests in the photoperiod-extension method (PEM), as well as at (c) the plot level in the conventional plot trials. Below the heat maps, are indicated the number post-flowering hot days with maxima exceeding 30°C from 0 to 500°Cd after flowering for each environment, grouped by heat environment types are indicated by dashed, coloured boxes (HET1, green; HET2, orange; HET3, red). All environments were fully irrigated except TOS19 (framed in brown), which experienced a mild post-flowering water stress. Genetic correlations for total grain weight are presented in Figure S2.

In the conventional plot trials, genotype ranking for IGW varied widely across environments, both between sowings and sites (Fig. 7c). Correlations between non-stressed environments (HET1) were relatively high (*r* of 0.4-0.78). However, correlations between HET2 environments were highly variable or even negative (*r* of -0.15 to 0.42). Similarly, HET3 trials GAT19s2 and TOS19s2 had a correlation of 0.10.

Grain yield from PEM quadrat harvests for irrigated trials were also positively correlated. The correlations ranged from 0.18 to 0.72 within HET1, from 0.33 to 0.63 within HET2 (Fig. S2b, supplementary data). In contrast, for conventional plots, correlations for grain yield were either poor or negative for most trials, irrespective to their heat environment types (Fig. 7c). For instance, maximum correlations within HET1, HET2 and HET3 were 0.3, 0.21 and -0.36, respectively.

Genotype rankings across methods were also compared. Strong positive relationships (*r*^2^ from 0.65 to 0.97 depending on the environment considered) were observed between the IGW from PEM trials with (i) individual spike harvest and (ii) quadrat harvest under the fully irrigated conditions (Fig. 8a). In contrast, correlations for IGW between PEM quadrat harvest and conventional plot trials under irrigated conditions were poorer in most trials (*r*^2^ from 0.19 to 0.57) (Fig. 8b.

**Figure 8:**
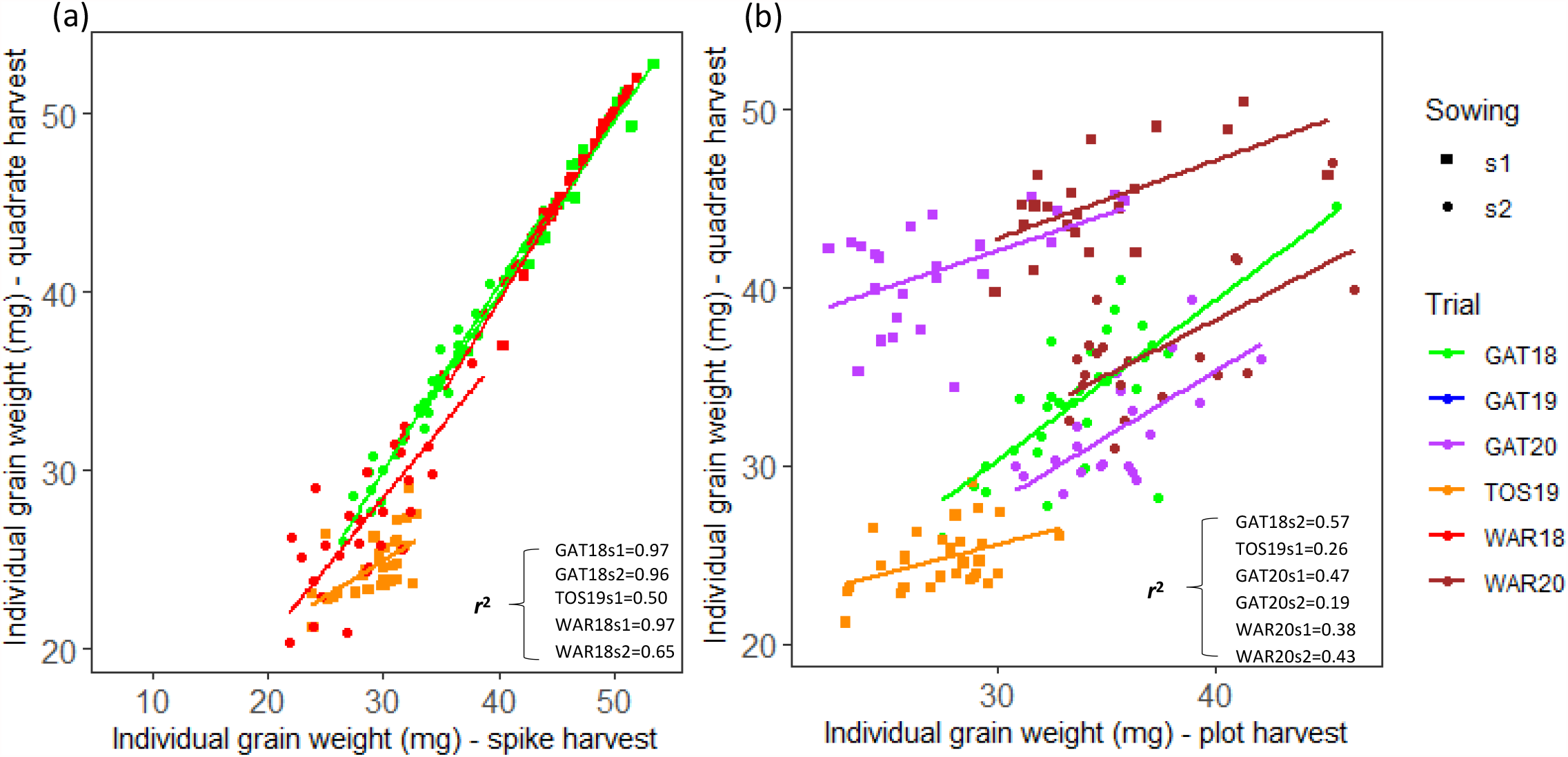
Correlations for individual grain weight of all studied genotypes between either (a) data collected from individually tagged stems and tagged quadrats at the matched development stage with the photoperiod-extension method (PEM), or between (b) the PEM with quadrat tagging and harvests and conventional plots. Each data point represents the genotypic mean value of four independent replicates. Correlations for total grain weight are presented in Figure S1.

### The photoperiod-extension method allowed genotypes to be discriminate according to their performance in different type of heat environments

Performance of wheat genotypes in terms of IGW and total grain weight were analysed using a principal component analysis (PCA, Fig. 9). The first two principle components (PC) combined explained more than 50% variance across environments in both IGW and grain yield for all the tested methods, except for grain yield in conventional field plots (Fig. 9).

**Figure 9:**
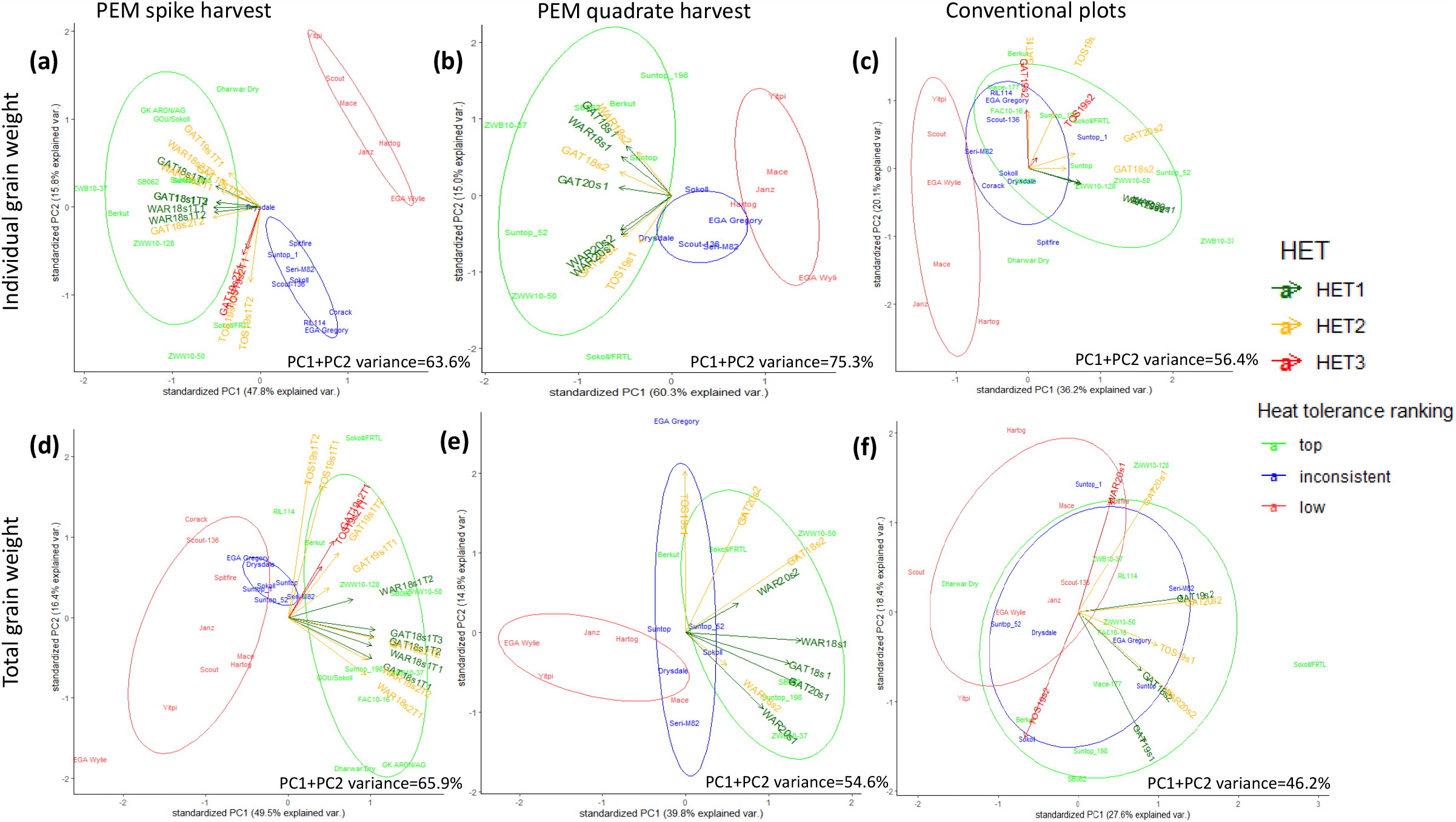
Principal component analysis biplots of (a-c) individual grain weight (IGW) and (d) grain weight per spike or (e-f) grain yield of studied wheat genotypes for the PEM spike harvests (a, d), PEM quadrat harvests (b, e) and conventional machine harvested plot trials (c, f). Environments corresponded to combinations of sowing dates, sites and years (b, c, e, f) together with tagging events for single-spike harvests (a, d). Principal component loadings (arrows) were coloured and grouped based on heat environment type; HET1, green; HET2, orange; HET3, red. Genotypes were grouped for similarity in their performance for either IGW (a-c) or total grain weight (d-f) in PEM spike harvests, i.e. green, top performing genotypes in HET1 and HET2, blue, genotypes with inconsistent / poor performance, particularly across HET1 and HET2; red, genotypes with poor performance across most tested environments. Eclipse colours correspond to the performance groups for genotypes.

For PEM spike harvests, strong positive correlations were observed between fully-irrigated HET1 and HET2 environments as indicated by the small angles between the vectors in the biplots (Fig. 9a & d). These correlations were stronger within HETs for IGW (Fig. 9a) than for total grain weight per spike (Fig. 9d). In contrast, a wider angle between vectors for HET3 and HET1-HET2 suggested weaker correlations between these environments (Fig. 9 a & d). Drought affected TOS19s1, which also experienced moderate post-flowering heat stress (HET2) better correlated with HET3 environments severely stressed by heat both for IGW and total grain weight per spike.

The projection of a genotype onto an environmental axis reflects the performance of that genotype in that environment. Genotypes were grouped based on their performance in environments from the PEM spike harvests. Biplots separated genotypes into three distinct groups for IGW and three partly differentiated groups for grain weight per spike (Fig. 9a & d). Top performing genotypes in HET1 and HET2 such as ZWB10-37 (in green in Fig. 9) project above the origin on axes corresponding to HET1 and HET2. In contrast, poorly performing genotypes such as Yitpi and EGA Wylie (in red in Fig. 9) project below the origin on the same axes. The top and poor performing genotypes identified with the PEM spike harvests were consistent with those identified from the PEM quadrat harvests data (Fig. 9b & e).

For the conventional plots, heat environment types (HET) were less clearly differentiated, particularly for grain yield, indicating less power to distinguish between HETs (Fig. 9c & f). Genotype rankings were also changed compared to ranking with the PEM. For instance, Suntop_1 ranked among the top performing genotypes for IGW in HET1 and HET2 in the conventional plot trials (Fig. 9c) but it ranked with the intermediate to poor group with the PEM (Fig. 9a). For grain yield in conventional plot trials, genotype performance was even more difficult to differentiate. For example, top performing genotypes in the PEM (in green) were not distinct from others in conventional plots being scattered almost randomly across the biplot (Fig. 9f).

## Discussion

### Impact of post-flowering heat events on individual grain weight and yield

Crop simulation studies identified post-flowering heat as the major determinant of wheat productivity in the Australian wheatbelt for current and future climates (Ababaei and Chenu, 2020; Collins and Chenu, 2021). Data from the current study demonstrates experimentally how heat intensity during the grain filling period is a major determinant for IGW and grain yield in irrigated field conditions (Fig. 6), agreeing with previous reports for crops grown under rainfed field conditions (Gebeyehou et al., 1982; Gerard et al., 2020; Telfer et al., 2018; Thistlethwaite et al., 2015).

In wheat, temperatures above 30°C negatively impact different processes during the grain filling period (Porter and Gawith, 1999; Girousse et al., 2021). In the tested conditions, under full irrigation, wheat genotypes did not experience any significant loss in IGW or grain yield in response to 0-4 hot days (maximum daily temperature > 30°C) during grain filling (HET1, Fig. 6a). These heat events were either very brief (1-2 h only in total from 0 to 450°Cd after flowering) or the heat event occurred late in development when grain filling was already well advanced i.e. > 450°Cd after flowering (Fig. 2). By contrast, in environments with 5 or more days with maximum temperatures >30°C between flowering and 500°C after flowering (≥4 cumulated hours of temperature >30°C; i.e. HET2 & HET3), wheat genotypes were substantially impacted. A significant and strong effect of the timing of heat stress has also been recorded for wheat IGW during early to mid grain filling in both controlled (Stone and Nicolas, 1996) and field conditions (Thistlethwaite et al., 2015). In the field conditions tested in the current study, each additional hot day during grain filling reduced IGW by 1.5 mg (Fig. 6a). This significant (*p*<0.001) reduction in IGW was also found in plants sown at single date but exposed to a range of natural heat events. For example, in GAT19s1, stems tagged one week apart (taggings 1 & 2) experienced different levels of heat stress, with stems from tagging 2 being subjected to four additional days of post-flowering heat and producing grain 14% smaller than stems from tagging 1 (Fig. 6a, Table S3). Developmental-phase-specific effects of heat shocks on IGW and grain yield were observed in this study, particularly for moderate stress in HET2 (data not shown). This highlights the importance of screening for heat tolerance at matched developmental phases. High heat sensitivity during early grain filling has also been reported by Talukder et al. (2013), who observed a single hot day (maximum temperature > 35°C) during this period can significantly reduce grain weight of wheat crops.

### A new method to screen for heat tolerance at matched development stages

During both the reproductive and the grain filling phases, sensitivity of developing grains to heat events can change over a period as short as a few days (Stone and Nicolas, 1995; Chenu and Oudin, 2019). Since field-based techniques for screening heat tolerant germplasm generally rely serial sowings, it is hard to optimise sowing time given the unpredictable timing of heat events. Field-based screening for heat tolerance using this conventional method is further complicated when genotypes with varying maturity types are tested together. These genotypes are likely to be at different developmental stages when natural heat events occur. Thus, heat escape by being at a less sensitive stage can confound comparisons of heat tolerance, *per se*.

We developed a new method using supplemental lights that allows screening of wheat genotypes at matched developmental phases. Photoperiod was extended to 20 h at one end of test rows of plots. The light intensity diminishes with the square of distance (Niinemets and Keenan, 2012), generating a gradient of flowering times along the length of the test rows (Fig. 1 & 4). The range of flowering times within a single row of individual genotype, allowed comparison of the performance of genotypes with varying maturity types at matched developmental stages. The method was tested for sowing dates from late May to mid-September. The impact of extended photoperiod on flowering was associated with the natural photoperiod during vegetative crop growth (*r*^2^ = 0.32; Fig. 5). A wide gap in flowering along the rows, of up to 8.8 days when averaged across all tested genotypes, was recorded when planting under short photoperiod (<10.50 h). This gap narrowed by 3.5 days with each increasing hour in photoperiod at later sowing dates (Fig. 5), so that all genotypes flowered within only 4.5 days of each other in the latest sowing dates tested. A wider flowering-time gradient is interesting as it allows; (i) multiple taggings within a single time of sowing, which may increase the probability of being able to screen for heat at a particular stage, (ii) more flexibility for operators to visit the trial during the appropriate window and tag all genotypes at a matched development stage, and/or (iii) a wider range of maturity types to be considered. Despite this additional flexibility, the robust genotype rankings generated with the PEM suggest that a single tagging is likely sufficient for reliable screening.

In the conditions tested, sowing dates for screening post-flowering heat tolerance between early July and late August provided the best discrimination. Sowings during this period were typically associated with moderate post-flowering heat stress (HET2) and a relatively wide flowering gradient for tagging stems or quadrats at a matched development stage. With earlier sowings, very few heat events occurred. In contrast, later sowings had a narrower flowering gradient and a greater risk of exposure to a high number of severe heat events which can increase heat damage to a level where variation between genotypes is reduced. Late sowings are also usually more prone to pre-flowering heat (e.g. TOS19s2 and GAT19s2), thus resulting in confounding effects, with IGW varying due to both a decrease in grain number and direct effects of post-flowering heat stress. Optimum sowing dates to screen heat stress obviously depend on the targeted stress (e.g. with or without pre-flowering heat). They also depend on the test location that impacts both wheat phenology and the frequency of heat events. Crop models can help to identify locations and sowing windows for efficient screening of heat tolerance in particular target environments (e.g. Chauhan et al., 2017; Chenu et al., 2017; Collins and Chenu, 2021).

### The PEM method allows reliable ranking of wheat genotypes under varying environments

With the PEM, strong correlations among trials for either IGW or grain yield (Fig. 7a & b, S2a & b) indicated a stable ranking of tested genotypes particularly within the environments receiving a similar degree of heat (Supplementary tables S4, S5 & S7). For instance, for the PEM with individual-spike harvests, the respective mean *r* correlations within fully-irrigated HET1, HET2 and HET3 environments were 0.80, 0.59 and 0.46 for IGW; and 0.75, 0.54 and 0.57 for total grain weight per spike. In contrast, these correlations were typically much weaker in conventional plots with *r* averages of 0.53, 0.11 and 0.1 for IGW, and 0.05, 0.12 and -0.36 for grain yield within HET1, HET2 and HET3, respectively (Fig. 7c, S2c, Supplementary tables S6 & S8). This indicates that in conventional plots, ranking of wheat genotypes was highly variable depending on environmental conditions, including for the targeted moderate grain-filling heat stress (HET2).

The PEM allowed the tested genotypes to be clustered into distinct groups based on their performance in these environments (Fig. 9a, b & d, e). In contrast, this genotype clustering was inconsistent with that found in conventional plots. The clustering in the conventional plots was also much less powerful at separating groups of genotypes as seen by the greater overlap of genotypes within each group (Fig. 9c & f). This indicates that the PEM ranked wheat genotypes more consistently across a wide range of heat-stressed environments compared to conventional plot trials.

### Implications for breeding

A high-throughput and accurate method for screening heat tolerance is necessary for sustaining food security under changing environments, particularly for projected hot and dry environments (Collins and Chenu, 2021). The PEM was tested using single rows (hand planted) and small plots (machine planted) with tagging of either individual stems or small quadrats at distances from the lights where development was matched at the flowering stage. The most reliable genotype ranking was achieved by tagging and harvesting individual spikes (Fig. 7a, 9a & d, S2a). However, strong positive relationships for either IGW (*r*^*2*^ from 0.65 to 0.97) or total grain weight (*r*^2^ from 0.34 to 0.97 across environments) between individual-spike and quadrat harvests were observed for the PEM (Fig. 8a & S1), suggesting that reliable screening may be done with the PEM for small plot with quadrat harvest. This was supported by the results, as genotype ranking with quadrat harvest was also robust (Fig. 7b, 9b & e, S2b) with strong genetic correlations between tested environments (average *r* of 0.51 in HET1 and 0.53 in HET2 for IGW, and 0.47 and 0.36 for grain yield). Using the PEM, with genotypes sown in small plots with machinery, and harvested from plot segments tagged at a matched developmental stage could have great potential to be scaled-up for large number of genotypes.

In the tested conditions, the PEM allowed effective screening for post-flowering heat stress (HET2), which is the main type of heat stress targeted for Australian production regions in current and projected climates (Collins and Chenu, 2021). Robust screening was also performed for non-stressed environments (HET1) and severely stressed environments (HET3) that happened to be affected by both pre- and post-flowering heat in the tested conditions. Different genotype rankings were observed between heat environment types HET2 and HET3, highlighting the importance of screening in the relevant target HET. The occurrence of water stress also strongly impacted genotype ranking (e.g. Fig. 2 and S2), suggesting that heat and drought adaptations are at least partly regulated by different processes. This highlights the importance of understanding the physiology and genetics associated to heat tolerance, drought tolerance and their interaction.

The PEM described here offers an opportunity to select for heat tolerant wheat genotypes more reliably than can be done conventionally. This method could be adjusted and deployed in other regions and other crops, with sowing dates adapted to the targeted heat environment type.

## Conclusions

A new field-based method was developed and tested to screen wheat genotypes for post-flowering heat tolerance under natural heat events. In this method, supplemental light was used to manipulate crop phenology in a way that allowed genotypes with varying phenology to be tested at closely matched developmental stage when a heat event occurred. IGW of wheat genotypes tested in this study was highly sensitive to the number of hot days during early-to-mid grain filling, particularly in the most relevant heat environment for Australian wheat crops (HET2). With the PEM, strong genetic correlations were found between irrigated environments of similar heat stress, with respective *r* correlations within HET1, HET2 and HET3 averaging 0.8, 0.59 and 0.46 for IGW; and 0.75, 0.54 and 0.57 for total grain weight. By contrast these correlations were substantially weaker for conventional yield plots (average *r* of 53, 0.11 and 0.1 for IGW; and 0.05, 0.12 and -0.36 for grain yield). Clustering of genotypes across tested environments with a PCA further highlighted the ability of the PEM to differentiate genotypes based on their performance in similar heat-stress environments. In contrast, ranking substantially changed between environments with moderate (HET2) and severe (HET3) heat stress. Similarly, genotype rankings greatly varied between fully-irrigated HET2 environments and HET2 environments subjected to a post-flowering water stress.

With increasing frequency of post-flowering heat stress affecting wheat growing regions, the photoperiod-extension method promises to improve the efficiency of heat tolerance field screening, particularly when comparing genotypes of different maturity types.

## Acknowledgments

The research was made possible thanks to the support of The University of Queensland and the Queensland Government, Department of Agriculture and Fisheries. Najeeb Ullah was supported by a Queensland Government, Advance Queensland Fellowship. We acknowledge the assistance of Ian Broad (Department of Agriculture and Fisheries Queensland) with operating of weather stations across the studied sites, Brian Collins (James Cook University) for data analysis and Thaís Helena Godoy Sanches for sample processing and data collection.

