## supplementary data for "A robust field-based method to screen heat tolerance in wheat"

**Table S1**: Characteristics of wheat genotypes used in the study

| Genotypes | Type | Pedigree | Characteristics | Reference | Year tested |
|---|---|---|---|---|---|
| Berkut | Cultivar | Irena/Baviacora-M-92//Pastor | Heat tolerance | (Thistlethwaite et al., 2020) | 2018, 2019, 2020 |
| Corack | Cultivar | MACHETE/84W129-504*2/5/VICAM S 71//CIANO F 67/SIETE CERROS/ 3/KAL/BB/4/TM56 | Australian Premium White (APW) variety | (GRDC, 2018) | 2018, 2019 |
| Dharwar Dry | Cultivar | DWR39/C306//HD2189 | Drought tolerant, stay-green phenotype | (Manschadi et al., 2008) | 2018, 2019 |
| Drysdale | Cultivar | Hartog*3/Quarrion | Superior transpiration efficiency | (Fletcher et al., 2018; Lobell et al., 2015; Richard et al., 2015) | 2018, 2019, 2020 |
| EGA Gregory | Cultivar | Pelsart/2*Batavia | Long-season elite cultivar with broad adaptation | | 2018, 2019, 2020 |
| EGA Wylie | Cultivar | INIA F 66 / GAMUT// COOK/3/JUPATECO F 73/TR 59 | Disease resistance | (Zheng et al., 2014) | 2018, 2019, 2020 |
| FAC10-16 | Elite breeding line | 10CB-F/W234 | Disease resistance | (Dinglasan et al., 2016) | 2018, 2019 |
| Fang | Cultivar | ANNUELLO/2*STYLET | Heat tolerance | (Skylas et al., 2002) | 2018 |
| GK ARON/AG | Elite breeding line | GK ARON/AG SECO 7846//2180 /4/2*MILAN/KAUZ//PRINIA/3/BAV92 | Heat tolerance | (Thistlethwaite et al., 2020) | 2018, 2019 |
| GOU/Sokoll | Elite breeding line | GOUBARA-1/2*Sokoll | Heat tolerance | (Thistlethwaite et al., 2020) | 2018, 2019, 2020 |
| Hartog | Cultivar | Vicam 71//Ciano 's'/Siete Cerros/3/Kalyansona/Bluebird | Drought sensitive, senescent phenotype | (Christopher et al., 2008; Richard et al., 2015) | 2018, 2019, 2020 |
| Janz | Cultivar | 3-AG-3/4*CONDOR//COOK | Drought sensitive, senescent phenotype | Christopher et al., 2008 | 2018, 2019, 2020 |
| Mace | Cultivar | Wyalkatchem/Stylet//Wyalkatchem | Benchmark cultivar for yield across southern Australian cropping regions | (GRDC, 2018) | 2018, 2019, 2020 |
| Mace-177 | Elite breeding line | Mace/Seri-M82 | Drought adaptation | (Christopher et al., 2021) | 2018, 2019 |
| PBW343 | Cultivar | NORD-DESPREZ/VG-1944//KALYANSONA//BLUEBIRD/3/ YACO(SIB)/4/VEERY-5 | Heat tolerant | (Thistlethwaite et al., 2020) | 2018 |
| RIL114 | Elite breeding line | UQ01484/RSY10//H45 | Pre-harvest sprouting tolerance | (Hickey et al., 2009) | 2018, 2019 |
| SB003 | Elite breeding line | Seri-M82/Babax | Heat sensitive, low stem water soluble carbohydrates | (Dreccer et al., 2009; Olivares-Villegas et al., 2007; Rattey et al., 2009; Ullah and Chenu, 2019) | 2019, 2020 |
| SB062 | Elite breeding line | Seri-M82/Babax | Heat tolerant, high stem water soluble carbohydrates | (Dreccer et al., 2009; Olivares-Villegas et al., 2007; Rattey et al., 2009; Ullah and Chenu, 2019) | 2018, 2019, 2020 |
| Scout | Cultivar | Sunstate/QH71-6//Yitpi | Elite cultivar with broad adaptation | | 2018, 2019, 2020 |
| Scout-136 | Elite breeding line | Scout/RIL114 | Drought sensitive, senescent phenotype | (Christopher et al., 2021) | 2018, 2019, 2020 |
| Seri-M82 | Elite breeding line | Kavkaz/4/Saric F 70///Lerma Rojo 64A/Inia F66//Inia F66/Yecora F70/5/II-26992 | Dense root system, stay-green phenotype | (Christopher et al., 2008; Olivares-Villegas et al., 2007) | 2018, 2019, 2020 |
| Sokoll | Cultivar | Pastor/3/Altar84/AE.SQ (TR.TA)//OPATA-M-85 | Heat tolerant | (Thistlethwaite et al., 2020) | 2018, 2019, 2020 |
| Sokoll//FRTL | Elite breeding line | SOKOLL//FRTL/2*PIFED | Heat tolerant | (Thistlethwaite et al., 2020) | 2018, 2019, 2020 |
| Spitfire | Cultivar | Drysdale/Kukri | Heat tolerant, high grain protein content | | 2018, 2019 |
| SSrT17 | Elite breeding line | Double backcross for the TIN gene into the free-tillering Silverstar background | Low tillering (*tin1* allele), big grains | (Mitchell et al., 2008) | 2019 |
| SSrW35 | Elite breeding line | Double backcross for the TIN gene into the free-tillering Silverstar background | High tillering (wild type allele) | (Mitchell et al., 2008) | 2019 |
| Suntop | Cultivar | Sunco/2*Pastor//SUN436E | Elite cultivar with broad adaptation; superior transpiration efficienc | (Collins et al., 2021; Richard et al., 2015) | 2018, 2019, 2020 |
| Suntop_1 | Elite breeding line | Suntop /Dharwar Dry | Drought tolerant, stay-green phenotype | (Christopher et al., 2021) | 2018, 2019 |
| Suntop _198 | Elite breeding line | Suntop/SB062 | Drought tolerant, stay-green phenotype | (Christopher et al., 2021) | 2018, 2019, 2020 |

| | | | | | |
|---|---|---|---|---|---|
| Suntop _52 | Elite breeding line | Suntop /Dharwar Dry | Drought tolerant, stay-green phenotype | (Christopher et al., 2021) | 2018, 2019, 2020 |
| WH 542 | Cultivar | BJY/JUP//URES | Heat tolerant | (Thistlethwaite et al., 2020) | 2018 |
| Yitpi | Cultivar | Condor/Gabo | Long coleoptile, very susceptible to yellow spot | | 2018, 2019, 2020 |
| ZWB10-37 | Elite breeding line | Tacupeto F2001/Brambling//Kiritati | High yield in CIMMYT-Australia-ICARDA Germplasm Evaluation | (Dinglasan et al., 2016) | 2018, 2019, 2020 |
| ZWW10-128 | Elite breeding line | ESDA/KKTS | High yield in CIMMYT-Australia-ICARDA Germplasm Evaluation | (Dinglasan et al., 2016) | 2018, 2019 |
| ZWW10-50 | Elite breeding line | Onix/4/Milan/Kauz//Prinia/3/BAV92 | High yield in the CIMMYT-Australia-ICARDA Germplasm Evaluation | (Dinglasan et al., 2016) | 2018, 2019, 2020 |

**Table S2**: Environmental conditions across trials and taggings in trials with the photoperiod-extension method (PEM). Data were averaged from sowing to maturity. The first, second and third taggings are referred to as 'T1', 'T2' and 'T3', respectively.

| Trial | Period | Tagging | Average daily temperature (°C) | Average daily max temp. (°C) | Mean day-time VPD (kPa) |
|---|---|---|---|---|---|
| GAT18s1 | Pre-flowering | First (T1) | 14.8 | 24.2 | 0.90 |
| | | Second (T2) | 15.1 | 24.5 | 0.95 |
| | | Third (T3) | 15.3 | 24.6 | 0.93 |
| | Post-flowering | First (T1) | 19.6 | 26.3 | 0.74 |
| | | Second (T2) | 20.2 | 26.5 | 0.76 |
| | | Third (T3) | 20.6 | 26.6 | 0.80 |
| GAT18s2 | Pre-flowering | First (T1) | 18.6 | 25.6 | 0.75 |
| | | Second (T2) | 20.2 | 25.9 | 0.82 |
| | Post-flowering | First (T1) | 23.3 | 31.4 | 1.38 |
| | | Second (T2) | 23.6 | 32.1 | 1.41 |
| WAR18s1 | Pre-flowering | First (T1) | 13.2 | 22.4 | 0.91 |
| | | Second (T2) | 13.5 | 22.5 | 0.95 |
| | Post-flowering | First (T1) | 18.8 | 25.6 | 0.84 |
| | | Second (T2) | 19.8 | 26.4 | 0.85 |
| WAR18s2 | Pre-flowering | First (T1) | 18.4 | 25.3 | 0.83 |
| | | Second (T2) | 18.9 | 25.6 | 0.93 |
| | Post-flowering | First (T1) | 22.1 | 30.3 | 1.52 |
| | | Second (T2) | 21.8 | 30.8 | 1.55 |
| GAT19s1 | Pre-flowering | First (T1) | 16.0 | 22.3 | 0.95 |
| | | Second (T2) | 16.1 | 22.6 | 0.96 |
| | Post-flowering | First (T1) | 18.0 | 28.7 | 1.40 |
| | | Second (T2) | 18.4 | 29.3 | 1.44 |
| GAT19s2 | Pre-flowering | First (T1) | 18.8 | 28.2 | 1.44 |
| | Post-flowering | First (T1) | 21.8 | 33.8 | 2.23 |
| TOS19s1 | Pre-flowering | First (T1) | 14.6 | 24.1 | 0.89 |
| | | Second (T2) | 15.0 | 24.5 | 0.90 |
| | | Third (T3) | 15.2 | 24.8 | 0.91 |
| | Post-flowering | First (T1) | 21.5 | 30.2 | 1.26 |
| | | Second (T2) | 21.8 | 30.8 | 1.31 |
| | | Third (T3) | 22.0 | 31.2 | 1.36 |
| TOS19s2 | Pre-flowering | First (T1) | 19.7 | 28.7 | 1.17 |
| | Post-flowering | First (T1) | 26.6 | 36.7 | 2.27 |
| GAT20s1 | Pre-flowering | First (T1) | 15.0 | 21.9 | 0.64 |
| | Post-flowering | First (T1) | 17.9 | 26.8 | 1.08 |
| GAT20s2 | Pre-flowering | First (T1) | 17.7 | 26.7 | 1.02 |
| | Post-flowering | First (T1) | 22.2 | 31.2 | 1.35 |
| WAR20s1 | Pre-flowering | First (T1) | 11.3 | 19.7 | 0.58 |
| | Post-flowering | First (T1) | 18.2 | 26.8 | 0.79 |
| WAR20s2 | Pre-flowering | First (T1) | 15.0 | 23.7 | 0.79 |
| | | Second (T2) | 15.3 | 24 | 0.81 |
| | Post-flowering | First (T1) | 22.1 | 29.5 | 1.40 |
| | | Second (T2) | 22.3 | 29.9 | 1.42 |

**Table S3**: Means and *P* values for grain number (GN), individual grain weight (IGW, mg) and total grain weight (GW, g) of wheat genotypes tested with (i) the photoperiod-extension method (PEM) with tagging and harvesting of individual spikes, (ii) the PEM with tagging and harvesting of quadrats, and (iii) conventional plots across trials. IGW and GN values correspond to mean of all studied genotypes and four replications. Grain number and grain weight data are presented as (i) per $m^2$ for PEM quadrat harvests and conventional plots, and (ii) per spike for spike harvest. *, **, *** indicate significant differences at P<0.05, P<0.01, P<0.001, respectively, and NS corresponds to 'non-significant'. Trial identifiers are as described in Table 1.

| **Harvesting** | **Site** | **Gatton** | | | | | | | | | **Tosari** | | | **Warwick** | | | | | |
|---|---|---|---|---|---|---|---|---|---|---|---|---|---|---|---|---|---|---|---|
| | **Trial** | **GAT18** | | | **GAT19** | | | **GAT20** | | | **TOS19** | | | **WAR18** | | | **WAR20** | | |
| | Trait | IGW | GN | GW | IGW | GN | GW | IGW | GN | GW | IGW | GN | GW | IGW | GN | GW | IGW | GN | GW |
| **Individual spikes (PEM)** | Sowing1 | 43.1 | 33.2 | 1.43 | 37.7 | 31 | 1.16 | | | | 27.2 | 40.3 | 1.10 | 46. | 32.4 | 1.50 | | | |
| | Sowing2 | 34.6 | 32.6 | 1.13 | 12.8 | 28 | 0.35 | | | | 17.8 | 29 | 0.52 | 28.1 | 32.3 | 0.91 | | | |
| **P values** | Sowing | *** | * | *** | *** | *** | ** | | | | *** | *** | *** | *** | NS | *** | | | |
| | Genotype | *** | *** | *** | *** | *** | *** | | | | *** | *** | *** | *** | *** | *** | | | |
| | Tagging | *** | *** | *** | *** | *** | NS | | | | *** | *** | *** | * | * | *** | | | |
| | Sowing×Genotype | *** | *** | *** | NS | NS | NS | | | | ** | ** | * | *** | *** | *** | | | |
| | Sowing×Tagging | *** | *** | NS | NS | NS | NS | | | | NS | NS | NS | *** | *** | *** | | | |
| | Genotype×Tagging | *** | *** | *** | | | | | | | | | | ** | ** | *** | | | |
| **Quadrats (PEM)** | Sowing1 | 45.5 | 12556 | 513 | | | | 41.6 | 6760 | 317 | 24.6 | 2289 | 316 | 45.1 | 9229 | 286 | 45.0 | 10292 | 437 |
| | Sowing2 | 33.9 | 12863 | 398 | | | | 31.9 | 7035 | 241 | | | | 27.5 | 8302 | 208 | 36.1 | 9029 | 348 |
| **P values** | Sowing | *** | * | *** | | | | *** | NS | *** | | | | *** | *** | *** | *** | *** | *** |
| | Genotype | *** | *** | *** | | | | *** | NS | *** | | | | *** | *** | *** | *** | * | *** |
| | Sowing×Genotype | *** | *** | *** | | | | NS | NS | NS | | | | *** | *** | *** | *** | * | ** |
| **Conventional plots** | Sowing1 | | | | 36.8 | 5254 | 192 | 28.2 | 9112 | 249 | 27.5 | 9945 | 272 | | | | 37.7 | 12838 | 478 |
| | Sowing2 | 33.8 | 6476 | 220 | 25.0 | 3966 | 99 | 35.5 | 4382 | 162 | 24.2 | 2589 | 62 | | | | 33.2 | 10290 | 362 |
| **P values** | Sowing | | | | *** | *** | *** | *** | *** | *** | *** | *** | *** | | | | *** | *** | *** |
| | Genotype | | | | *** | *** | *** | *** | ** | *** | *** | *** | *** | | | | *** | *** | *** |
| | Sowing×Genotype | | | | *** | *** | *** | *** | NS | NS | *** | *** | *** | | | | *** | *** | *** |

**Table S4**: Individual grain weight (IGW, mg) of all studied genotypes in trials with the photoperiod-extension method (PEM) with tagging and harvesting of individual spikes. Genotypes are ordered based on the average individual grain weight across trials with low post-flowering heat stress, i.e. heat environment type 1 (HET1). Colours indicate the rankings of the genotypes within each trial, going from red (lowest IGW) to green (highest IGW).

| **Heat environment type** | **HET1** | | | | | **HET2** | | | | | | | | | **HET3** | |
|---|---|---|---|---|---|---|---|---|---|---|---|---|---|---|---|---|
| Genotypes | GAT18 s1T1 | WAR18 s1T1 | WAR18 s1T2 | GAT18 s1T2 | GAT18 s1T3 | GAT19 s1T2 | WAR18s2T1 | WAR18 s2T2 | GAT18 s2T1 | GAT18 s2T2 | GAT19 s1T1 | TOS19 s1T1 | TOS19 s1T2 | TOS19 s1T3 | GAT19 s2T1 | TOS19 s2T1 |
| GK ARON/AG | 53.4 | 51.1 | 47.9 | 47.4 | 46.7 | 42.4 | 31.9 | 31.9 | | 36.5 | 42.4 | 28.3 | 25.1 | 26.4 | 9.9 | 17.7 |
| PBW343 | 51.6 | 51.2 | 47.0 | 48.7 | | | 27.8 | 30.9 | 35.4 | 34.9 | | | | | | |
| ZWB10-37 | 51.5 | 49.8 | 52.4 | 45.5 | 48.9 | 41.9 | 38.7 | 34.1 | 45.9 | 43.8 | 41.9 | 29.7 | 24.7 | 25.0 | 16.1 | 18.6 |
| ZWW10-128 | 50.9 | 49.5 | 50.6 | 44.8 | 47.1 | 41.9 | 31.8 | 28.7 | 40.3 | 37.9 | 41.9 | 32.6 | 26.4 | 27.2 | 14.2 | 17.4 |
| Berkut | 50.8 | 51.9 | 54.2 | 49.1 | 42.2 | 45.1 | 32.3 | 30.3 | 40.4 | 38.1 | 45.1 | 31 | 27 | 27.7 | 12.7 | 18.9 |
| WH 542 | 50.6 | 49.8 | 43.0 | 48.7 | 42.1 | | 30.1 | 17.3 | 35.1 | 33 | | | | | 16.2 | 22.7 |
| Dharwar Dry | 50.4 | 44.8 | 46.6 | 42.9 | 43.0 | 42.6 | 31 | 27.7 | 38.2 | 35.6 | 36.6 | 25 | 22.6 | | 13 | 18.5 |
| GOU/Sokoll | 50.3 | 48.3 | 49.8 | 47.0 | 45.1 | 43.4 | 33.8 | 31.6 | | 36.5 | 43.4 | 30.9 | 25.3 | 25.7 | 10 | 16 |
| SB062 | 49.6 | 50.7 | 50.9 | 44.4 | 44.6 | 45.3 | 34.3 | 29.8 | 41.0 | 39.2 | 45.3 | 30.1 | 26.5 | 28.9 | 12.9 | 17.4 |
| Suntop_198 | 47.3 | 50 | 49.9 | 44.7 | 46.3 | 40.3 | 32.2 | 29.5 | 39.3 | 36.1 | 40.3 | 28.7 | 26.1 | 26.4 | 11.1 | 17.6 |
| ZWW10-50 | 46.8 | 49.9 | | 42.3 | 46.5 | 41 | 29.9 | 27.7 | | 36.4 | 41.0 | 31.3 | 29.3 | 23.3 | 15.2 | 20 |
| FAC10-16 | 46.6 | 48.9 | | 44.8 | 46.5 | 41.7 | 26 | 26.6 | | 34.3 | 41.7 | 27.7 | 24.7 | 26.7 | 14.7 | 17.3 |
| Suntop | 46.3 | 47.3 | 47.5 | 43.0 | | 44.7 | 31.5 | 31.1 | 35.4 | 34.2 | 44.7 | 31.6 | 26.4 | 26.6 | 10.6 | 18.3 |
| Drysdale | 46.2 | 47.4 | 47.4 | 42.6 | 41.4 | 40.9 | 26.9 | | 38.8 | 31 | 40.9 | 30.7 | 25.1 | 26.5 | 11.6 | 17.4 |
| Suntop_52 | 45.8 | 49.1 | 49.4 | 41.0 | 41.7 | 41.5 | 37.6 | 36.0 | 38.5 | 36.5 | 41.5 | 30.8 | 25.4 | 26.6 | 12.2 | 18 |
| Corack | 45.3 | 45.1 | 48.0 | 33.6 | 38.5 | 43.2 | 23.9 | 21.2 | | 29.8 | 43.2 | 31.3 | 25.5 | 26.1 | 11.2 | 20.6 |
| Scout-136 | 45 | 46.2 | 46.7 | 38.7 | 43.8 | 39.6 | 26.2 | 25.3 | 34.8 | 30 | 39.6 | 31.9 | 27.3 | 25.5 | 12.1 | 17.4 |
| Suntop_1 | 44.3 | 44.5 | 46.9 | 44.6 | 42.9 | 37.5 | 22.1 | 29.4 | 34.4 | 31.7 | 37.5 | 29.7 | 26.6 | 27.5 | 12.3 | 17.7 |
| Yitpi | 44.2 | 44.1 | 47.5 | 35.7 | 41.0 | 41.4 | 28.5 | 29.7 | | 29 | 41.4 | 25.3 | 23.3 | 22.6 | 8.7 | 15.6 |
| Spitfire | 44 | 46.4 | 43.2 | 37.3 | 39.2 | 39.2 | 27 | 27.5 | | 33.6 | 39.2 | 29.3 | 25.5 | 23.3 | 12.7 | 19.1 |
| Hartog | 44 | 41.5 | 39.6 | 39.2 | 40.0 | 35.9 | 25 | 24.7 | 32.5 | 27.4 | 35.9 | 23.8 | 23.2 | | 13.9 | 16.4 |
| Sokoll/FRTL | 43.7 | 43.7 | 47.9 | 41.2 | 42.7 | 42.2 | 32 | 26.1 | 36.4 | 34.7 | 42.2 | 32 | 27.7 | 31.1 | 15.7 | 19.4 |
| Seri-M82 | 43.6 | 43.4 | 44.7 | 43.0 | 41.2 | 40.8 | 24.6 | 23.4 | 30.8 | 33 | 40.8 | 31.3 | 26.7 | 26.9 | 14.8 | 15.4 |
| Sokoll | 43.3 | 42.9 | 44.4 | 41.0 | 39.8 | 38.5 | 27.9 | 27.3 | 36.0 | 33.3 | 38.5 | 31.2 | 27.2 | 27.3 | 12.8 | 17.3 |
| Mace | 42.6 | 42.2 | 42.4 | 37.2 | 39.7 | 42 | 24.1 | 26.2 | | 29.2 | 42.0 | 25.8 | 24 | 20.4 | 8.6 | 18.3 |
| Scout | 42.4 | 43.9 | 44.7 | 35.7 | 40.4 | 37.9 | 24 | 23.8 | 34.8 | 31.1 | 43.9 | 26 | 23.4 | | 9.9 | 16.1 |
| Janz | 41.2 | 40.5 | 41.9 | 35.5 | 36.7 | 42.2 | 28.7 | 24.6 | 33.8 | 28.9 | 42.2 | 24 | 24.3 | 22.8 | 13.7 | 18.5 |
| RIL114 | 41 | 44 | 43.6 | 37.4 | 39.4 | 43.1 | 28.6 | 24.3 | 37.7 | 33.8 | 43.1 | 32.9 | 26.6 | 24.7 | 13.6 | 18.1 |
| EGA Gregory | 40.6 | 43 | 46.0 | 36.2 | 37.8 | 39.6 | 23 | 26.6 | | 33.9 | 44.6 | 32.2 | 28.3 | 27.8 | 11.4 | 18.3 |
| EGA Wylie | 37.3 | 40.4 | 41.2 | 33.1 | 35.4 | 38.2 | 29.8 | 26.1 | 32.6 | 28.8 | 38.2 | 28.4 | 27.1 | 25.1 | 9.1 | 14.9 |
| Fang | 35.1 | 35.3 | 37.6 | 29.8 | 30.9 | 36.3 | 21.9 | 24.0 | 28.0 | 26.6 | 36.3 | | | | 7.1 | |
| Mace-177 | | 45.3 | | 42.5 | 45.7 | 43.4 | | 25.6 | | 33.6 | 43.4 | 30.1 | 24.1 | 25.6 | 12 | 16.2 |
| SB003 | | | | | | 37.3 | | | | | 37.3 | 30.1 | 24.3 | 26.2 | 12.9 | 17.7 |
| SsrT1 | | | | | | 39.3 | | | | | 39.3 | 30.1 | 26.8 | 29.1 | 9.1 | 17.3 |
| SsrW35 | | | | | | 39.5 | | | | | 39.5 | 29.8 | 26.3 | 27.2 | 13.6 | |

**Table S5**: Individual grain weight (IGW, mg) of all studied genotypes in trials with the photoperiod-extension method (PEM) with tagging and harvesting of quadrats. Genotypes are ordered based on the average individual grain weight across trials with low post-flowering heat stress, i.e. heat environment type 1 (HET1). Colours indicate the rankings of the genotypes within each trial, going from red (lowest IGW) to green (highest IGW).

| **Heat environment type** | **HET1** | | | | | **HET2** | | | |
|---|---|---|---|---|---|---|---|---|---|
| Genotypes | GAT20s1 | WAR20s1 | WAR20s2 | GAT18s1 | WAR18s1 | GAT18s2 | GAT20s2 | TOS19s1 | WAR18s2 |
| Suntop_52 | 45.2 | 46.3 | 47.0 | 45.3 | 49.4 | 37.0 | 39.3 | 24.6 | 27.7 |
| ZWW10-50 | 45.1 | 50.4 | 41.5 | 47.1 | 49.9 | 36.4 | 36.7 | 27.2 | 21.3 |
| ZWB10-37 | 44.9 | 49.0 | 39.9 | 49.2 | 49.9 | 44.6 | 36.0 | 24.9 | 31.4 |
| ZWW10-128 | 44.3 | | | 51.1 | 49.5 | 38.8 | | 23.7 | 20.9 |
| SB062 | 44.1 | 45.3 | 36.8 | 49.6 | 50.7 | 40.4 | 33.1 | 24.5 | 25.2 |
| Sokoll | 43.4 | 42.0 | 36.7 | 43.3 | 42.9 | 33.3 | 29.6 | 23.9 | 25.8 |
| EGA Wylie | 42.5 | 41.0 | 32.5 | 36.7 | 37.0 | 29.9 | 29.4 | 24.4 | 24.4 |
| Suntop | 42.5 | 44.5 | 36.1 | 47.0 | 47.3 | 34.2 | 31.8 | 25.6 | 20.3 |
| Sokoll/FRTL | 42.4 | 48.9 | 41.7 | 43.7 | 43.7 | 34.7 | 35.2 | 27.3 | 31.9 |
| Corack | 42.3 | | | 44.9 | 44.8 | 28.2 | | 26.1 | 31.4 |
| Janz | 42.2 | 43.1 | 35.1 | 41.2 | 40.5 | 28.9 | 30.3 | 21.2 | 25.8 |
| Berkut | 41.8 | 42.0 | 36.3 | 50.8 | 52.0 | 37.6 | 30.1 | 26.0 | 24.6 |
| Suntop_198 | 41.7 | 44.1 | 35.2 | 47.9 | 50.1 | 36.1 | 29.2 | 23.3 | 29.0 |
| SB003 | 41.2 | 48.3 | 35.1 | | | | 30.0 | 23.9 | 25.9 |
| Hartog | 40.7 | 43.5 | 33.9 | 44.0 | 41.5 | 28.5 | 32.2 | 23.1 | 25.9 |
| EGA Gregory | 40.5 | 44.5 | 34.5 | 40.6 | 43.0 | 33.3 | 29.6 | 25.0 | 24.4 |
| Yitpi | 39.9 | 39.7 | 32.5 | 44.2 | 44.1 | 27.7 | 28.4 | 22.8 | |
| GK ARON/AG | 39.7 | | | 52.8 | 51.1 | 36.5 | | 24.1 | 29.8 |
| Mace | 38.3 | 43.5 | 30.9 | 41.6 | 40.9 | 30.8 | 30.0 | 22.9 | 23.8 |
| Mace-177 | 37.6 | 46.3 | 36.0 | | 45.3 | 32.4 | 30.0 | 23.5 | 25.2 |
| Seri-M82 | 37.2 | 44.6 | 35.9 | 42.8 | 43.4 | 33.5 | 31.1 | 24.7 | 22.9 |
| Scout-136 | 37.0 | 44.6 | 34.5 | 45.0 | 46.2 | 30.0 | 33.6 | 25.8 | 27.3 |
| Scout | 35.3 | | | 42.4 | 44.4 | 30.9 | | 23.2 | 25.6 |
| Drysdale | 34.4 | 45.5 | 39.4 | 46.2 | 47.4 | 33.6 | 34.2 | 23.7 | 27.5 |
| Dharwar Dry | | | | 50.7 | 44.6 | 34.3 | | 26.4 | |
| FAC10-16 | | | | 45.2 | 48.9 | 35.0 | | 23.1 | |
| Fang | | | | 35.1 | 35.3 | 26.0 | | | 31.1 |
| GOU/Sokoll | | | | 50.6 | 48.3 | 37.9 | | 23.9 | 26.3 |
| PBW343 | | | | 49.3 | 51.2 | 36.8 | | | 29.5 |
| RIL114 | | | | 41.0 | 44.0 | 33.8 | | 27.6 | 36.0 |
| Spitfire | | | | 43.0 | 46.4 | 33.9 | | 26.3 | |
| SsrT1 | | | | | | | | 25.7 | 29.9 |
| SsrW35 | | | | | | | | 23.5 | 38.9 |
| Suntop_1 | | | | 44.3 | 44.2 | 31.7 | | 25.3 | 32.5 |
| WH542 | | | | 50.6 | 49.8 | | | | 27.7 |

**Table S6**: Individual grain weight (IGW, mg) of all studied genotypes in conventional plots. Genotypes are ordered based on the average individual grain weight across trials with low post-flowering heat stress, i.e. heat environment type 1 (HET1). Colours indicate the rankings of the genotypes within each trial, going from red (lowest IGW) to green (highest IGW).

| **Heat environment type** | **HET1** | | | **HET2** | | | | **HET3** | |
|---|---|---|---|---|---|---|---|---|---|
| Genotypes | GAT20s1 | WAR20s1 | WAR20s2 | GAT18s2 | GAT19s1 | GAT20s2 | TOS19s1 | GAT19s2 | TOS19s2 |
| ZWB10-37 | 35.8 | 46.4 | 37.4 | 45.6 | 33.3 | 42.2 | 25.8 | 24.9 | 26.3 |
| Suntop_52 | 35.5 | 45.4 | 45.2 | 32.5 | 37.8 | 38.9 | 29.2 | 25.3 | 23.4 |
| ZWW10-128 | 32.8 | 44.9 | 32.0 | 35.4 | 36.7 | 35.7 | 29.0 | 23.9 | 24.2 |
| Suntop_1 | 32.9 | 41.9 | 35.5 | 32.0 | 37.4 | 38.0 | 27.8 | 27.0 | 24.6 |
| Suntop_198 | 24.6 | 41.5 | 33.7 | 36.8 | 36.1 | 36.3 | 29.6 | 26.9 | 25.7 |
| ZWW10-50 | 31.6 | 41.1 | 41.3 | 37.9 | 40.2 | 38.1 | 28.1 | 22.2 | 24.1 |
| Sokoll/FRTL | 29.2 | 41.0 | 40.7 | 35.0 | 38.8 | 35.5 | 30.1 | 26.0 | 23.4 |
| Dharwar Dry | 29.3 | 40.1 | 34.1 | 36.4 | 36.6 | 32.6 | 24.3 | 22.4 | 23.9 |
| SB003 | 27.3 | 40.1 | 34.4 | | 38.9 | 34.7 | 30.1 | 26.0 | 25.0 |
| Suntop | 32.5 | 39.3 | 35.6 | 33.8 | 35.4 | 37.0 | 29.3 | 25.9 | 22.8 |
| FAC10-16 | 27.9 | 38.9 | 26.1 | 34.6 | 41.5 | 35.9 | 27.0 | 24.7 | 23.8 |
| Hartog | 29.4 | 37.6 | 33.2 | 29.5 | 32.2 | 33.7 | 23.3 | 21.8 | 24.9 |
| Spitfire | 29.9 | 36.6 | 35.7 | 32.5 | 34.2 | 38.6 | 26.3 | 23.1 | 22.2 |
| GK ARON/AG | 25.7 | 36.3 | 31.8 | 34.3 | 37.1 | 35.0 | | 23.9 | 24.2 |
| Seri-M82 | 25.2 | 36.0 | 31.7 | 33.5 | 37.8 | 33.7 | 25.8 | 26.0 | 24.7 |
| EGA Wylie | 23.3 | 35.8 | 31.7 | 34.0 | 37.1 | 31.2 | 24.7 | 25.9 | 22.6 |
| Scout-136 | 24.7 | 35.6 | 31.2 | 29.5 | 36.5 | 39.3 | 27.5 | 26.2 | 25.9 |
| Mace | 25.4 | 35.4 | 31.3 | 31.9 | 36.3 | 30.9 | 23.2 | 21.9 | 25.4 |
| Corack | 23.8 | 35.4 | 31.9 | 37.4 | 36.3 | 35.7 | | 25.3 | 24.1 |
| Sokoll | 26.0 | 34.8 | 36.4 | 33.1 | 36.9 | 33.8 | 29.1 | 23.4 | 24.0 |
| WH 542 | 24.0 | 34.8 | 32.4 | 37.0 | | 35.5 | 27.6 | | 24.9 |
| Drysdale | 28.1 | 34.6 | 36.3 | 32.9 | 34.7 | 35.6 | 27.4 | 25.5 | 24.5 |
| Berkut | 24.4 | 34.6 | 34.2 | 35.0 | 40.7 | 34.8 | 32.9 | 24.9 | 23.5 |
| Scout | 23.6 | 34.4 | 28.2 | 30.8 | 38.0 | 34.3 | 25.7 | 26.5 | 22.9 |
| SB062 | 27.0 | 34.2 | 33.4 | 35.6 | 35.3 | 36.2 | 28.5 | 23.6 | 25.0 |
| Janz | 22.3 | 34.0 | 33.6 | 28.9 | 32.7 | 32.6 | 23.1 | 22.1 | 22.6 |
| EGA Gregory | 27.2 | 33.9 | 32.3 | 32.3 | 35.6 | 36.2 | 28.9 | 28.1 | 25.7 |
| Mace-177 | 26.5 | 33.7 | 31.9 | 34.1 | 38.5 | 36.0 | 28.8 | 27.1 | 23.8 |
| RIL114 | 25.3 | 33.4 | 31.5 | 31.0 | 37.1 | 37.1 | 29.1 | 25.9 | 28.9 |
| Yitpi | 24.4 | 33.3 | 29.9 | 32.3 | 38.7 | 33.0 | 25.6 | 28.7 | 22.9 |
| Fang | | | | 27.5 | | | | | |
| GOU/Sokoll | | | | 36.6 | 36.1 | | 28.2 | 24.7 | 22.0 |
| PBW343 | | | | 37.1 | | | | | |
| SsrT1 | | | | | 36.7 | | 28.3 | 24.7 | 24.1 |
| SsrW35 | | | | | 37.3 | | | 26.6 | |

**Table S7**: Grain yield (g $m^{-2}$) of all studied genotypes in trials with the photoperiod-extension method (PEM) with the tagging and harvesting of quadrats. Genotypes are ordered based on the average individual grain weight across trials with low post-flowering heat stress, i.e. heat environment type 1 (HET1). Colours indicate the rankings of the genotypes within each trial, going from red (lowest IGW) to green (highest IGW).

| **Heat environment type** | **HET1** | | | | | **HET2** | | | |
|---|---|---|---|---|---|---|---|---|---|
| Genotype | GAT20s1 | WAR20s1 | WAR20s2 | GAT18s1 | WAR18s1 | GAT18s2 | GAT20s2 | TOS19s1 | WAR18s2 |
| ZWB10-37 | 424.3 | 450.9 | 332.6 | 541.2 | 495.8 | 440.0 | 282.8 | 254.2 | 292.5 |
| Mace-177 | 394.3 | 476.0 | 357.45 | | 440.4 | 388.6 | 232.2 | 294.4 | 177.2 |
| Suntop_198 | 363.1 | 458.9 | 342.1 | 604.8 | 463.1 | 505.5 | 264.0 | 242.4 | 278.0 |
| ZWW10-50 | 360.7 | 449.8 | 385.7 | 596.7 | 548.3 | 568.5 | 277.0 | 385.1 | 202.5 |
| Sokoll | 355.1 | 485.3 | 365.6 | 571.8 | 351.6 | 416.1 | 245.5 | 411.0 | 219.0 |
| Sokoll/FRTL | 343.2 | 471.9 | 426.9 | 497.8 | 366.9 | 446.9 | 366.0 | 320.1 | 219.6 |
| SB062 | 342.8 | 486.7 | 351.5 | 614.2 | 548.3 | 470.8 | 272.9 | 308.6 | 195.0 |
| Mace | 334.1 | 451.2 | 327.3 | 512.0 | 323.4 | 382.6 | 266.7 | 255.0 | 218.7 |
| Seri-M82 | 329.2 | 494.4 | 344.0 | 595.0 | 382.3 | 386.6 | 209.7 | 299.8 | 136.4 |
| SB003 | 326.1 | 446.1 | 326.3 | | | | 227.4 | 274.6 | |
| Drysdale | 324.3 | 462.2 | 326.1 | 504.6 | 393.0 | 391.1 | 269.4 | 221.6 | 139.4 |
| EGA Gregory | 316.3 | 391.3 | 301.5 | 501.6 | 361.1 | 478.7 | 278.6 | 533.2 | 188.2 |
| ZWW10-128 | 315.8 | | | 515.5 | 389.2 | 506.5 | | 231.3 | 214.2 |
| Scout-136 | 310.2 | 380.6 | | 512.4 | 359.2 | 372.8 | 220.9 | 321.7 | 175.0 |
| Suntop_52 | 305.5 | 464.4 | 349.1 | 457.0 | 333.8 | 510.2 | 282.9 | 256.9 | 274.6 |
| Berkut | 301.1 | 368.3 | 321.5 | 502.5 | 403.9 | 480.3 | 235.3 | 347.3 | 195.5 |
| Suntop | 298.2 | 526.0 | 380.1 | 367.6 | 335.1 | 365.5 | 220.8 | 376.5 | 232.2 |
| Yitpi | 292.6 | 387.7 | 320.9 | 304 | 305.8 | 219.8 | 180.6 | 293.1 | 214.0 |
| Scout | 286.9 | | | 429.1 | 369.6 | 323.3 | | 198.6 | 154.9 |
| Hartog | 279.9 | 388.1 | 415.0 | 514.0 | 358.9 | 330.7 | 192.2 | 295.8 | 176.9 |
| EGA Wylie | 276.1 | 341.9 | 280.2 | 345.6 | 251.0 | 309.3 | 170.0 | 316.1 | 202.4 |
| Janz | 275.5 | 400.6 | 302.2 | 440.8 | 308.2 | 318.6 | 228.9 | 302.9 | 214.3 |
| Corack | 261.0 | | | 541.8 | 318.6 | 243.6 | | 443.0 | 139.2 |
| GK ARON/AG | 247.3 | | | 735.4 | 536.7 | 485.7 | | 275.3 | 271.8 |
| Dharwar Dry | | | | 564.1 | 434.7 | 418.3 | | 454.6 | 180.5 |
| FAC10-16 | | | | 692.5 | 455.1 | 387.5 | | 246.2 | 223.2 |
| Fang | | | | 329.1 | 262.5 | 261.3 | | | 100.7 |
| GOU/Sokoll | | | | 580.2 | 373.1 | 412.3 | | 299.5 | 292.7 |
| PBW343 | | | | 429.4 | 411.4 | 446.9 | | | 228.6 |
| RIL114 | | | | 541.9 | 365.5 | 328.4 | | 399.8 | 201.1 |
| Spitfire | | | | 561.2 | 320.4 | 257.9 | | 376.0 | 212.9 |
| SsrT1 | | | | | | | | 383.5 | |
| SsrW35 | | | | | | | | 295.8 | |
| Suntop_1 | | | | 526.6 | 410.0 | 361.9 | | 386.7 | 212.5 |
| WH542 | | | | 435.2 | 389.5 | | | | |

**Table S8**: Grain yield (g $m^{-2}$) of all studied genotypes in conventional plots. Genotypes are ordered based on the average individual grain weight across trials with low post-flowering heat stress, i.e. heat environment type 1 (HET1). Colours indicate the rankings of the genotypes within each trial, going from red (lowest IGW) to green (highest IGW).

| **Heat environment type** | **HET1** | | | **TEH2** | | | | **HET3** | |
|---|---|---|---|---|---|---|---|---|---|
| Genotypes | GAT20s1 | WAR20s1 | WAR20s2 | GAT18s2 | GAT19s1 | GAT20s2 | TOS19s1 | GAT19s2 | TOS19s2 |
| ZWB10-37 | 248 | 508 | 357 | 300 | 165 | 201 | 256 | 76 | 56 |
| Seri-M82 | 294 | 503 | 398 | 273 | 190 | 156 | 324 | 115 | 55 |
| Scout-136 | 255 | 503 | 349 | 195 | 209 | 133 | 260 | 113 | 60 |
| Janz | 248 | 502 | 350 | 215 | 207 | 172 | 211 | 97 | 68 |
| Drysdale | 236 | 502 | 399 | 181 | 192 | 132 | 296 | 70 | 67 |
| Suntop_1 | 306 | 499 | 313 | 224 | 131 | 149 | 286 | 119 | 59 |
| Spitfire | 252 | 493 | 314 | 207 | 178 | 198 | 271 | 103 | 40 |
| SB003 | 241 | 491 | 386 | | 270 | 143 | 269 | 121 | 59 |
| ZWW10-128 | 355 | 489 | 403 | 215 | 182 | 156 | 246 | 116 | 33 |
| Hartog | 336 | 489 | 328 | 163 | 108 | 158 | 237 | 86 | 57 |
| Mace | 354 | 486 | 313 | 187 | 174 | 154 | 282 | 80 | 57 |
| EGA Gregory | 220 | 484 | 376 | 163 | 216 | 176 | 311 | 114 | 56 |
| RIL114 | 263 | 482 | 378 | 254 | 159 | 171 | 284 | 112 | 53 |
| Suntop | 314 | 481 | 441 | 225 | 243 | 190 | 257 | 89 | 69 |
| Mace-177 | 192 | 480 | 387 | 197 | 230 | 154 | 286 | 112 | 56 |
| Dharwar Dry | 258 | 479 | 280 | 159 | 149 | 131 | 274 | 77 | 77 |
| Sokoll/FRTL | 325 | 479 | 442 | 241 | 239 | 263 | 314 | 142 | 62 |
| SB062 | 229 | 479 | 408 | 295 | 227 | 150 | 284 | 71 | 81 |
| WH 542 | 238 | 474 | 383 | 321 | | 157 | 265 | | 62 |
| Yitpi | 185 | 473 | 341 | 167 | 206 | 131 | 221 | 89 | 78 |
| EGA Wylie | 214 | 468 | 361 | 166 | 170 | 142 | 238 | 100 | 60 |
| Scout | 199 | 467 | 241 | 198 | 150 | 132 | 245 | 54 | 57 |
| Suntop_52 | 157 | 465 | 236 | 214 | 188 | 141 | 269 | 108 | 53 |
| ZWW10-50 | 279 | 462 | 428 | 312 | 144 | 196 | 246 | 89 | 57 |
| Berkut | 249 | 460 | 377 | 176 | 193 | 159 | 274 | 80 | 84 |
| GK ARON/AG | 162 | 457 | 428 | 211 | 235 | 159 | | 119 | 65 |
| Suntop_198 | 176 | 450 | 396 | 295 | 213 | 140 | 260 | 115 | 56 |
| Sokoll | 163 | 446 | 398 | 273 | 160 | 133 | 285 | 83 | 63 |
| Corack | 218 | 442 | 400 | 136 | 182 | 188 | | 70 | 70 |
| FAC10-16 | 300 | 438 | 264 | 182 | 227 | 194 | 273 | 128 | 59 |
| Fang | | | | 187 | | | | | |
| GOU/Sokoll | | | | 212 | 192 | | 287 | 113 | 81 |
| PBW343 | | | | 289 | | | | | |
| SsrT1 | | | | | 174 | | 342 | 87 | 79 |
| SsrW35 | | | | | 272 | | | 124 | |

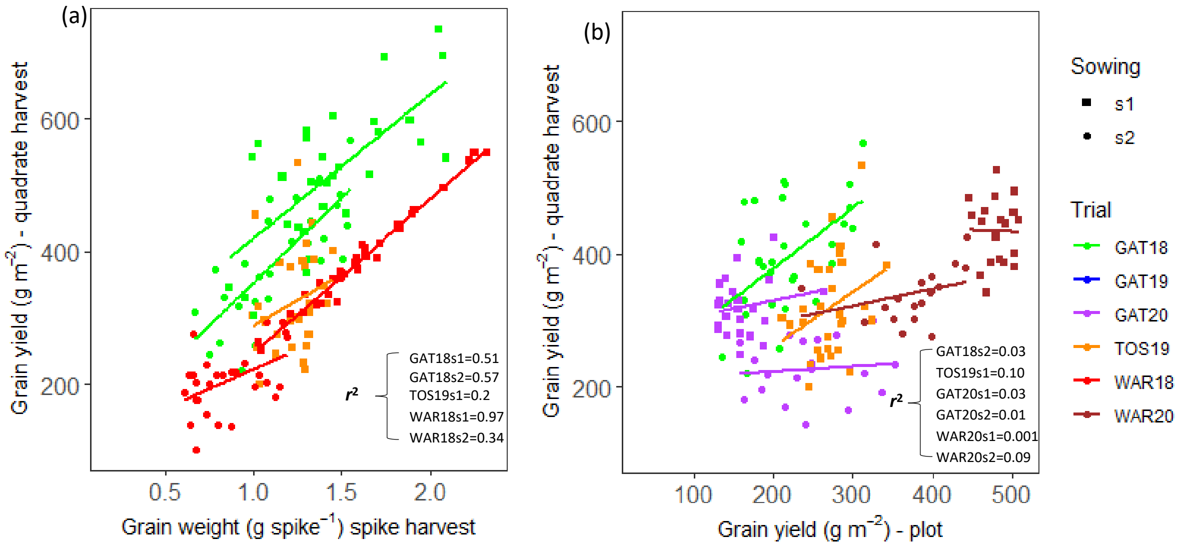


**Figure S1**: Correlations between quadrat harvests in photoperiod-extension trials and either individual-spike harvest (a, c) or plot trials (b, d) for individual grain weight (a, b) or total grain weight (c, d), i.e. grain weight per spike (c) or grain yield (d). Data correspond the mean of each of the studied genotypes (four independent replicates). TOS19 crops had supplementary irrigation and experienced a mild post-flowering water stress. Correlations for individual grain weight are presented in Figure 8.

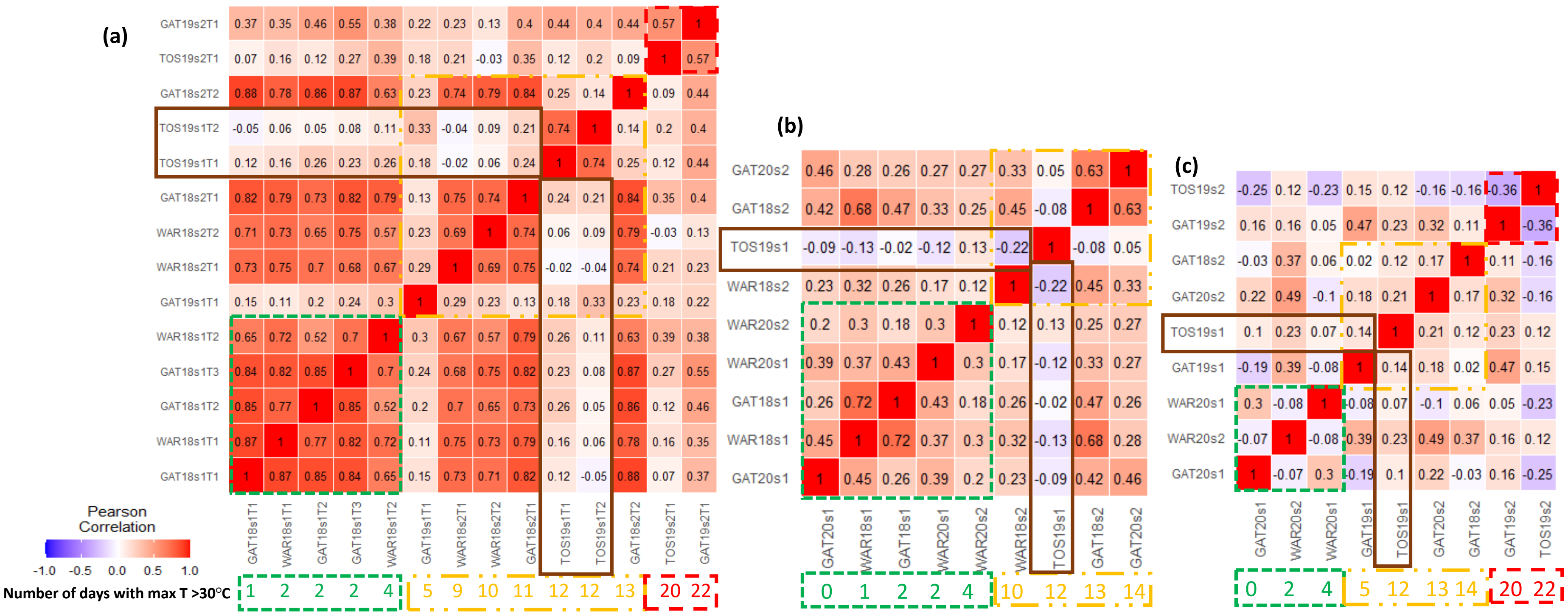

**Figure S2**: Genetic Pearson's correlation coefficients ($r$) of the genotype grain weight between pairs of studied environments with grain weight measured at (a) spike level as grain weight spike$^{-1}$, and at (b) crop level (g m$^{-2}$) with quadrat harvests in the photoperiod extension method (PEM), as well as at (c) the plot level in the conventional plot trials (g m$^{-2}$). Below the heat maps, in blue, are indicated the number post-flowering hot days (days with a maximum temperature above 30ºC between 0 and 500ºCd after flowering) for each environment, grouped by heat environment types (HET1, green; HET2, orange; HET3, red). All environments were fully irrigated except TOS19 (framed in brown), which experienced a mild post-flowering water stress. Correlations for individual grain weight are presented in Figure 7.
